# Deep-learning-based interpolation of longitudinal microbiome data powers biologically informative discovery

**DOI:** 10.1101/2025.02.17.638709

**Authors:** Yixiang Qu, Ruiqi Lyu, Duan Wang, Yifan Dai, Alistair Turcan, Shilin Yu, Jialiu Xie, Jeffrey Roach, Catherine Butler, Pew-Thian Yap, Hongtu Zhu, Stuart Dashper, Apoena Aguiar Ribeiro, Didong Li, Kimon Divaris, Di Wu

## Abstract

The human microbiome is a foundational and dynamic foundation for several health-related functions and disease processes. Advances in microbiome sequencing have enabled the characterization of microbial communities in several niches. Longitudinal microbiome studies further strive to discover clinically informative microbial community trajectories. However, these data are fraught with dropout events, high noise, and irregular sampling that limit and prevent the use of many available longitudinal analysis tools. To address these challenges, we introduce Bidirectional GRU-ODE-Bayes (BGOB), a deep learning framework developed for longitudinal microbiome interpolation. BGOB combines bidirectional information flow and ODE-based continuous modeling to jointly interpolate and smoothen trends across individual participants, providing uniform, denoised time intervals across patients. BGOB enables vastly improved performance in differential abundance testing and time-to-event analysis, and makes possible longitudinal analyses requiring uniformity, such as lead-lag detection and temporal clustering. After interpolation, previously low-powered datasets are able to broadly recapitulate known microbiology and elucidate interacting microbial communities. We highlight several associations between microbial taxa and disease, including novel species associated with Early Childhood Caries and disruption of key healthy gut microbiota in Inflammatory Bowel Disease. The BGOB package is publicly available at https://github.com/Rachel-Lyu/BGOB_n_test.

## 1 Introduction

Recent advances in sequencing technologies have enabled the taxonomic profiling and functional interrogation of entire microbiome communities, enabling novel insights for several clinical endpoints [1]. Longitudinal studies investigating microbiomes at multiple time points are well positioned to illuminate health-related aspects of development during childhood [2], disease progression [3], prediction of treatment outcomes [4] and dynamic monitoring of the environment [5]. Such data can capture temporal trajectories, microbe-microbe interactions, and time-varying microbe-disease associations [6], e.g., in childhood dental disease [7] and gut disease progression [8]. Microbiome research is faced with common challenges such as zero inflation, overdispersion, high dimensionality, and may entail normalized counts or compositional structures [9, 10, 11]. Critically, unlike traditional longitudinal data, microbiome samples in large cohorts are often collected at irregular intervals, and complete data at all time points are rare, even in well-designed studies (Figure 1), preventing analyses that require consistency in sample collection times between study participants. Furthermore, members of the microbial community often interact with each other, which undermines rigorous assumptions of a proper parametric distribution.

**Figure 1.**
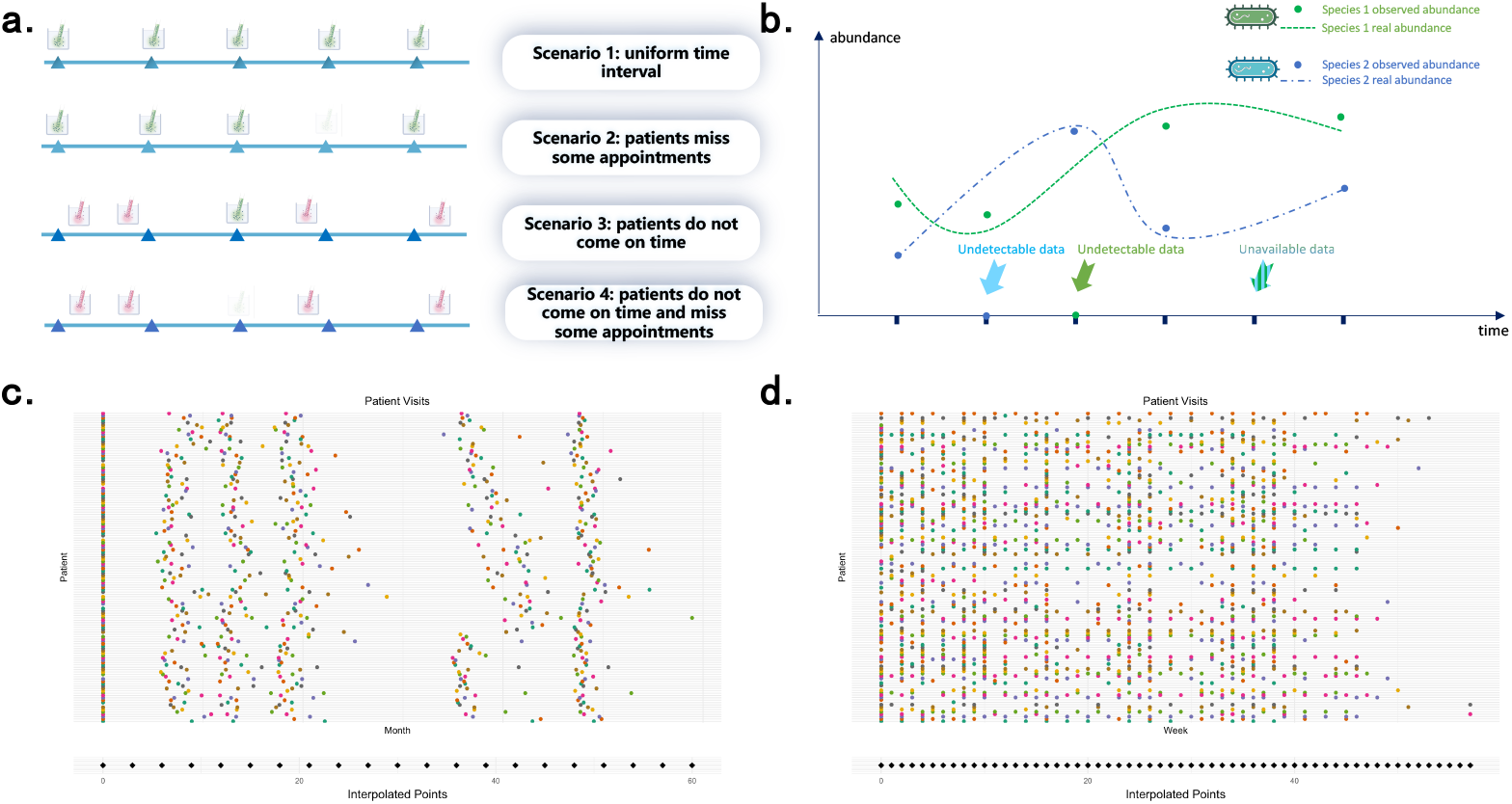
**(a)** Common scenarios that introduce irregular sampling in longitudinal microbiome data collection. Scenarios 2-4 generate inconsistent sample collection times among study participants. **(b)** Illustrative example of real vs observed trends for 2 species. Species 1 and 2 exhibit both observed zeroes (i.e., “undetectable” data) and missing time points (i.e., “unavailable” data). **(c)** Heterogenous sample collection dates across study participants in an ECC dataset, with BGOB’s uniform, interpolated intervals on the bottom. **(d)** Heterogenous sample collection dates across study participants in an IBD dataset, with BGOB’s uniform, interpolated intervals on the bottom.

Interpolation of longitudinal microbiome data can address irregular intervals and reconstruct temporal trajectories, enabling downstream data analysis. Longitudinal data interpolation, as a typical preprocessing option, has traditionally relied on statistical methods. Spline-based methods [12] can handle irregular sample collection [13, 14], but do not account for correlations between features and are not specifically designed for microbiome data. Most recent longitudinal microbiome models have not specifically addressed interpolation. The Temporal Gaussian Process Model for Compositional Data Analysis (TGP-CODA) [15], based on a Bayesian probabilistic model, leverages temporal correlations between time points between subjects to model microbial abundance, but is not explicitly designed for interpolation. LUMINATE (longitudinal microbiome inference and zero detection) [16] efficiently employs state-space models to accurately infer microbial relative abundances under zero inflation and compositional constraints, but cannot interpolate unseen time points. Recurrent Neural Networks (RNNs) are suitable for discrete longitudinal analysis without distributional assumptions [17], and Long Short-Term Memory (LSTM) [18], a type of RNN, has been applied to predict food allergies from microbiome data [19]. However, RNNs rely on fixed, discrete intervals for inference and struggle to represent varied sampling. Neural Ordinary Differential Equation (ODE)-based methods address this issue by enabling continuous-time modeling [20]. The GRU-ODE-Bayes (GOB) model [21] estimates the distributions at any time point by continuously modeling the hidden state. Although GOB has been successfully applied to prediction tasks, its potential for interpolation remains underexplored, and it has not been tailored to microbiome data.

Herein, we present the development and application of Bidirectional GRU-ODE-Bayes (BGOB), a deep learning model for reconstructing longitudinal microbiome data. We treat both the samples at absent time points (that is, “unavailable” data) and the zeroes in the observed samples (that is, “undetectable” data) as missing and aim to jointly infer their values. BGOB is specially designed for microbiome data, balancing the influence of zero-inflation and producing biologically relevant output. We further incorporate bidirectional information flow to enhance predictions of individual participants’ trends, especially at early time points. BGOB-interpolated data can subsequently be utilized to improve the performance of a wide range of downstream analyses, including but not limited to differential abundance (DA) testing, time-to-event analysis, lead-lag detection, and clustering. To the best of our knowledge, BGOB is the first deep learning method that can simultaneously reconstruct both unavailable and undetectable data. We perform extensive simulations showcasing BGOB’s superiority in reconstructing ground truth trends. We apply BGOB to oral microbiome data from a longitudinal cohort study examining early childhood caries (ECC) with irregular sampling intervals, and gut microbiome data examining Inflammatory Bowel Disease (IBD) with a wide range of dropout events. We find that the application of BGOB vastly increases the performance of downstream analyses, providing new insights into disease-relevant microbial communities, and underscoring the importance of interpolation in the data preprocessing step in longitudinal studies.

## 2 Results

### 2.1 BGOB Overview

Irregular sampling is a common challenge in longitudinal studies, often arising from missed visits or deviations from follow-up schedules (Figure 1). This leads to “unavailable” data, where samples are completely missing at specific time points. Furthermore, microbiome data inherently contain “undetectable” data, where certain species have zero abundance due to technical limitations or a true biological absence.

BGOB jointly infers the latent trends underlying each taxon in each individual participant, enabling interpolation of a smooth trend and imputation of undetectable zeroes at consistent intervals for every participant and taxon. This is done in the forward and backward directions simultaneously to ensure accuracy across all time points. We incorporate a modified Gated Recurrent Unit (GRU) to model the derivative of the hidden state with long-range dependencies across species and time. An ODE component enables continuous modeling to infer missing time points for any interval, effectively interpolating unavailable data. A “Bayesian-like loss function uses a sureness parameter to enable a confident prediction of the distribution of a species in both directions, along with a bespoke mask designed to prevent zero inflation (undetectable data) from overcontributing to the model. Lastly, we enforce biologically relevant output with a 5-layer neural network, squashing the values between 0 and 1. Further details are provided in the Methods and Supplementary Note.

Briefly, we discuss the conceptual differences between BGOB and other deep learning methods for longitudinal data analysis (Figure 2C). RNNs model sequential states iteratively; however, they require consistent time intervals to be effective, a rare occurrence in longitudinal microbiomics (Figure 1). GOB is an ODE-based interpolation method. ODE’s can continuously model hidden states using their derivatives to reconstruct entire trends, requiring no consistency of time intervals. GOB further incorporates a modified GRU to infer the derivative of the hidden states, allowing long-distance trend information to remain relevant. Both GOB and RNN’s operate in the forward direction only, negatively affecting predictions at the beginning of a trend. BGOB extends GOB in several ways, notably by jointly modeling trends both forward and backward to improve initial predictions, and ensuring output consistent with biology.

**Figure 2.**
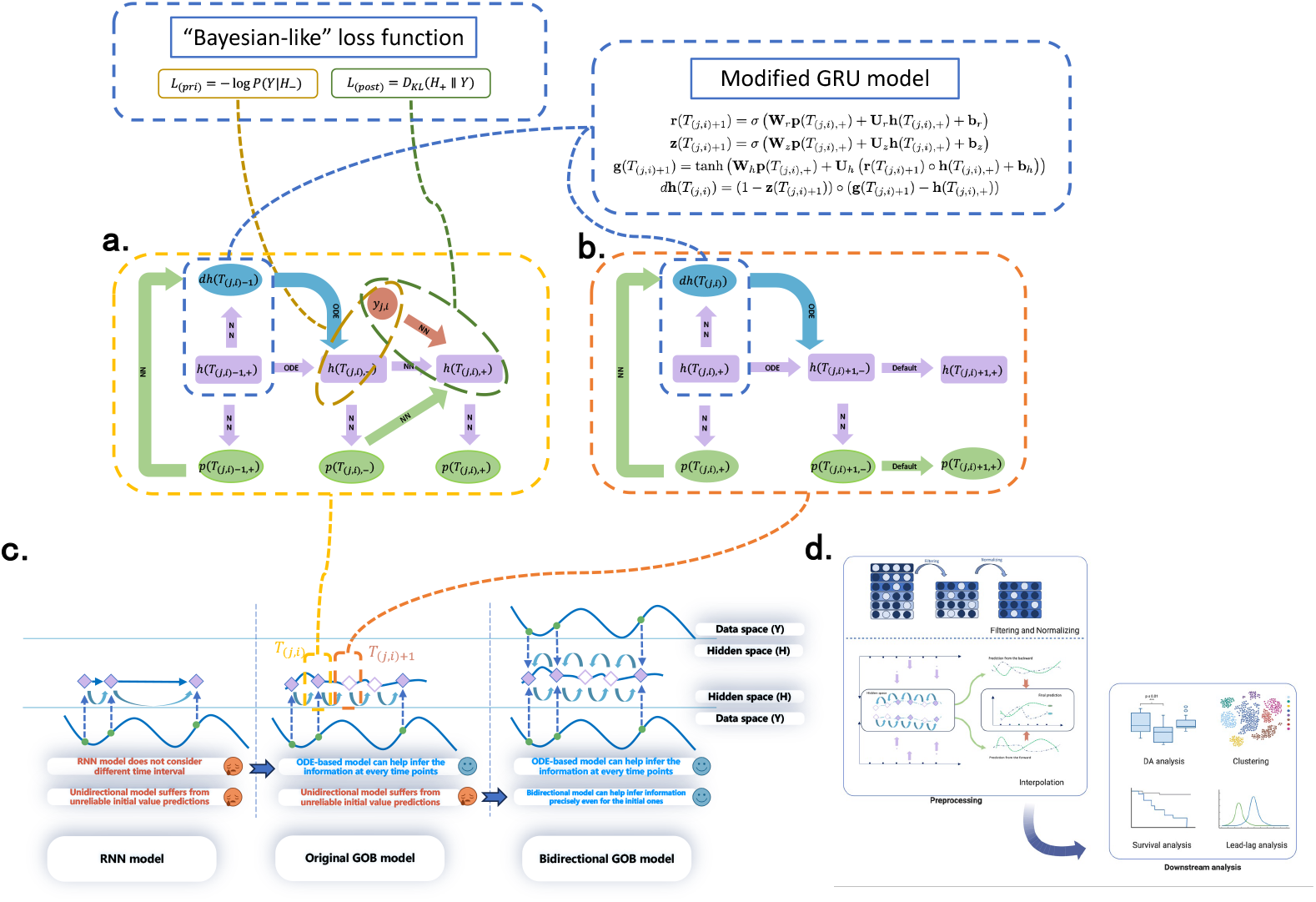
BGOB inputs longitudinal data and outputs interpolated and imputed time intervals for all individuals and species. **(a)** Information propagation in BGOB when observations are available nearby, mirrored in both the forwards and backwards direction. The derivative of the hidden state is inferred by a modified GRU and Bayesian-like loss function. The hidden state is propagated by an ordinary differential equation (ODE) with output post-processed by a neural network (NN). **(b)** Information propagation in BGOB when observations are not available nearby, jointly performed forwards and backwards. Default indicates equality between elements. **(c)** Comparison across model types: RNN (left), GOB (center), and BGOB (right). GOB, as one type of Neural ODE, extends RNNs by enabling continuous modeling. BGOB extends GOB by allowing for bidirectional information propagation. **(d)** BGOB analysis pipeline. Data are filtered and normalized, then interpolated with BGOB. Subsequently, longitudinal analyses can be performed, including but not limited to DA testing, time-to-event analysis, lead-lag detection, and clustering.

BGOB outputs interpolated and smoothed longitudinal data at specified intervals for each individual. The resulting data can then be applied to the wide variety of preexisting longitudinal analyses that require consistent time intervals. Here, we focus on 4 downstream analyses (Figure 2D). First, we perform differential abundance testing to identify species with higher or lower abundance in diseased individuals. Second, we performed a time-to-event analysis to discover microbes with disease-relevant trajectories, positing that they may either cause the disease or interact with a causal microbiome. Third, we perform lead-lag analysis, identifying leader and follower relationships across microbes, indicating potential microbe-microbe interactions, which may differ in the presence of disease. Fourth, we perform temporal clustering on a decomposed matrix of individuals and their time points, enabling the discovery of microbes that are correlated across time and may share interactions or environmental factors.

We perform comprehensive simulations, incorporating sparsity, missingness, and interactions among microbial taxa to compare various interpolation and imputation methods. Then, we analyze two real-world datasets. This includes an ECC dataset of 857 oral microbial species at up to 6 irregularly sampled time points for 132 children, 42 of whom developed ECC during the study. We also include an IBD dataset of 503 gut species and 130 individuals, with 1-26 visits each. We report results on both the original and interpolated data to demonstrate the necessity of interpolation as a preprocessing step.

### 2.2 Simulations Assessing Reconstruction Performance

We carry out and report two types of simulation. First, we conduct univariate simulations to capture independently fluctuating microbial taxa, measured in raw counts. Second, we carry out multivariate simulations with interacting microbial populations, measured in relative abundance (compositional). We compare interpolation accuracy and cluster purity scores across interpolation methods Cubic-Spline and GOB, as well as the accuracy of the interpolation method TGP-CODA [15] and the imputation method Luminate [16], which cannot interpolate data at unobserved time points.

We simulate univariate longitudinal data following a Zero-Inflated Log-Normal (ZILN) distribution (14) at 30 time points and 1000 individuals segregated into 5 distinct groups of equal size. The zero-inflation parameter represents the portion of the zeroes in the observed samples, or undetectable data. The microbiome community of each group is temporally modeled using a Black-Scholes (BS) [22] or Ornstein-Uhlenbeck (OU) [23] process (4.4). Each time point of each individual then has a possibility to be dropped out to simulate missed collection periods, the unavailable data. We vary both the zero inflation rate and the data dropout rate. Accuracy is measured with RMSE in raw count data (4.4.3). We observe a drastic increase in interpolation accuracy using deep learning methods over a simple cubic spline (Figure 3 A), particularly as the amount of information available decreases. BGOB provides a further improvement over GOB, particularly at the initial time points (Figure 3C) (mean improvement over GOB 28.07 ± 1.41% at the first 3 time points, 4.08 ± 1.35% at the remaining time points). When evaluating the purity of the cluster, where we group individuals according to their temporal trends, BGOB has the highest purity scores (Figure 3B). In particular, the score does not decrease considerably with less information, while Cubic Spline and GOB lose performance as data missingness increases.

**Figure 3.**
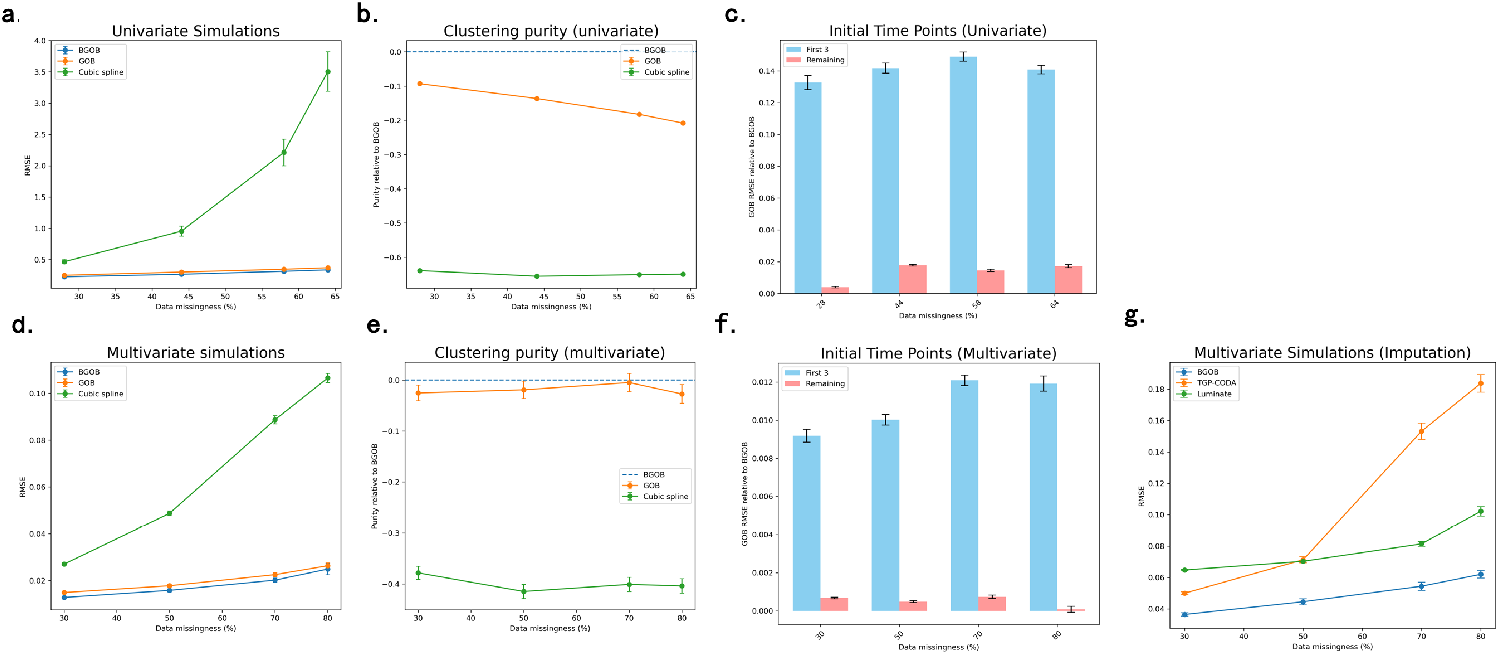
Simulation results assessing calibration and performance. Means and 95% confidence intervals of 20 simulation replicates are plotted for different levels of data missingness. **(a)** RMSE in univariate simulations, measured in raw counts. **(b)** Clustering purity scores (ranging from 0 to 1) relative to BGOB in univariate simulations. **(c)** RMSE in univariate simulations of GOB versus BGOB, highlighting performance at initial time points. Blue denotes the performance difference at the first 3 intervals, red denotes performance difference at the remaining 27 intervals. **(d)** RMSE in multivariate simulations, measured in relative abundance. **(e)** Clustering purity scores (ranging from 0 to 1) relative to BGOB in multivariate simulations. **(f)** RMSE in multivariate simulations of GOB versus BGOB, highlighting performance at initial time points. Blue denotes the performance difference at the first 3 intervals, red denotes performance difference at the remaining 27 intervals. **(g)** Imputation performance comparison between BGOB, TGP-CODA, and Luminate at only non-missing time intervals, measured in RMSE.

Multivariate data are additionally simulated following a ZILN distribution. 400 subjects are generated, divided into 5 equally sized groups, and observed at 30 time points each. Each subject contains 5 microbial species, again modeled with the BS and OU processes, with additional interspecies interactions based on the generalized Lotka-Volterra (gLV) model [24]. Microbial abundance depends both on the abundance of its previous time points, as well as on the abundance of other microbes (4.4.2). Unavailable and undetectable data missingness is again incorporated and varied. Here, RMSE is evaluated with relative abundance instead of raw counts (4.4.3). We again find that deep learning achieves higher accuracy and clustering purity scores compared to Cubic Spline and BGOB’s improved performance in both interpolation and clustering (Figures 3D, 3E). BGOB also achieves strong performance at the initial time points (Figure 3F) (mean improvement over GOB 32.29 ± 1.21% at the first 3 time points, 0.46 ± 2.33% at the remaining time points). Evaluating imputation accuracy alone (only at available data points) against TGP-CODA and Luminate, BGOB achieves the lowest RMSE (Figure 3G). BGOB’s accuracy also decreases much less as information decreases compared to TGP-CODA.

We conclude BGOB achieves higher performance across simulations and metrics compared to existing interpolation and imputation methods, and is well-suited for handling multiple sources of data missingness in both compositional and count data.

### 2.3 Dynamic Oral Microbe Development in ECC

We apply BGOB to a longitudinal ECC dataset from the VicGen cohort study [7]. Oral microbiome information is available for 132 individuals, 42 of whom developed ECC prior to their final visit. Data collection spans up to six time points, spaced from birth to roughly 48 months old (Figure 1C), with additional time-to-event data available for some individuals. We filter and aggregate 857 microbial species into 92 groups (4.5.1). For interpolation, we align continuous monthly values to integers of 3, and incrementally increase the overlapping intervals by 3 to ensure uniqueness. The maximum time point is set to 60 months, generating 21 discrete time points per subject.

We employ a regression-based differential abundance test (4.6) to identify species abundance-ECC associations (Figure 4A). We identified five significantly associated species (FDR*<* 0.1) after interpolation, while none are detected using the original data. These include *Porphyromonas pasteri, Veillonella dispar*, the *S. mutans* group, *Sneathia sanguinegens*, and *Leptotrichia shahii*. The signs of the coefficients for all these significant species do not change after interpolation, implying that BGOB does not introduce spurious associations. Most of the findings are broadly aligned with the known ecology, including higher abundances of *Veillonella sp*. and *Streptococcus mutans* (which belongs to the *S. mutans* group) being associated with disease, since their presence together results in greater acid production and increased demineralization compared with *S. mutans* alone [25, 26, 27, 28, 29, 30, 31, 32]. *L. shahii* has also been previously observed with higher abundance in ECC [33]. *P. pasteri* has a small reduction in caries-active individuals (coef = −2.01 × 10^−3^), consistent with recent studies [34] [35]. However, *P. pasteri* ‘s role in ECC is not well understood. *P. pasteri* is an obligately anaerobic, non-pigmented, non-spore-forming, non-motile, Gram-stain-negative rod—one of the 12 most abundant bacteria in the oral cavity, the second most abundant microbe in saliva [36]. Additionally, this species has been identified as part of the predominant commensal bacterial species found on the tongues of primary school children [37]. Notably, Mark Welch et al. [38] identified *Porphyromonas spp*., likely including *P. pasteri*, as a key member of health-associated “hedgehog” structures in supragingival plaque, which facilitate complex interactions with taxa such as *Leptotrichia, Streptococcus*, and *Corynebacterium*. Due to its prevalence in oral microbiomes, particularly those of children, it is plausible *P. pasteri* has a potential protective role in ECC. The negative association of *S. sanguinegens* with ECC in this study is intriguing, as this species, predominantly observed in vaginal microbiome disorders, is also present in oral cavities [39]. Notably, *S. sanguinegens* exhibits an unusual prevalence pattern, increasing in early childhood before declining over time, while being less prevalent in mothers [7]. Its unique dynamics and potential role in childhood oral health merit further investigation.

**Figure 4.**
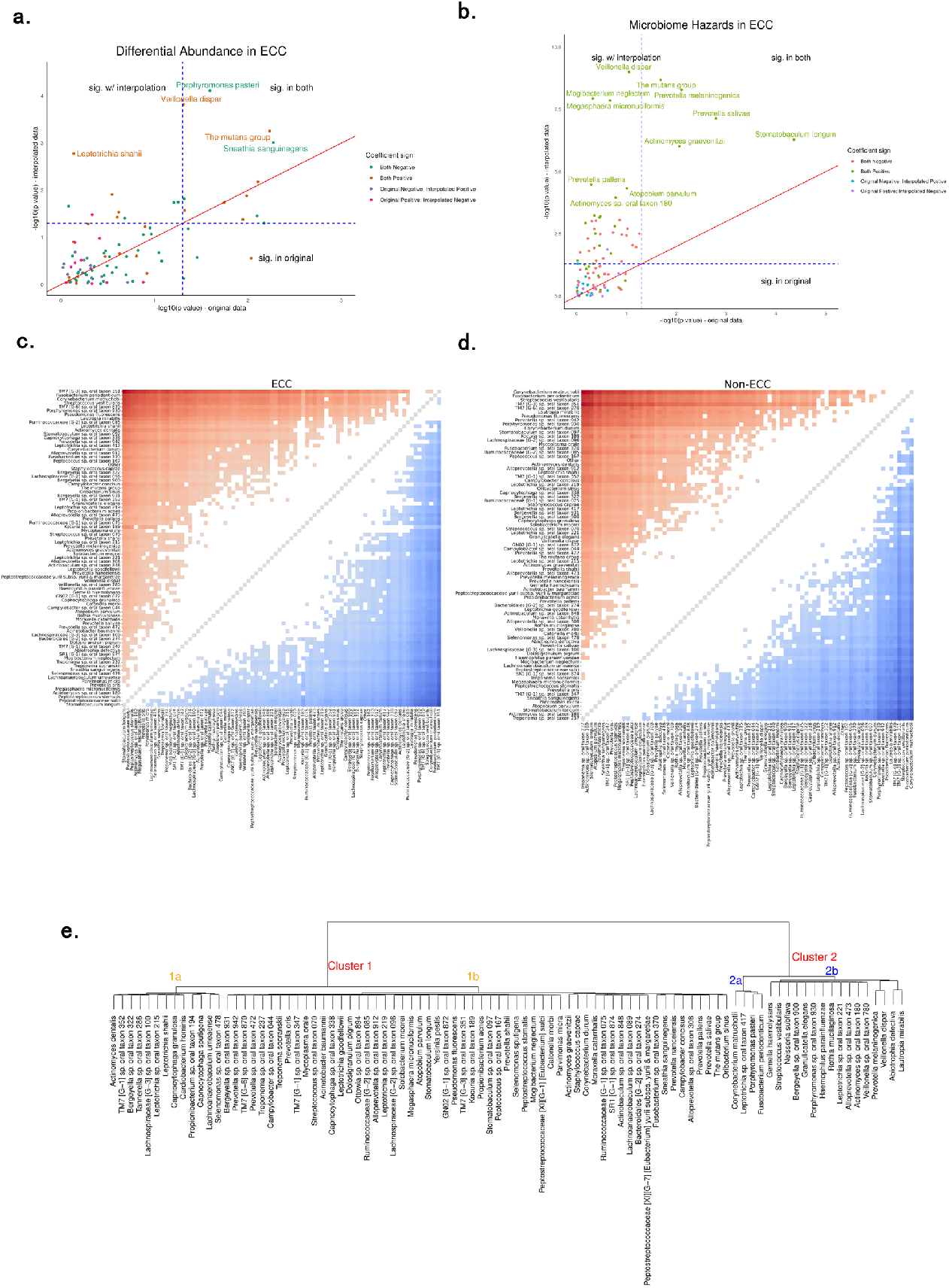
**(a)** Differential abundance results using the original data (x-axis) versus BGOB’s interpolated data (y-axis). Species with p-values less than 0.05 on either dataset are highlighted. Dot colors denote concordance between the abundance coefficients in the original and the interpolated data. **(b)** Time-to-event results using the original data (x-axis) versus BGOB’s interpolated data (y-axis). Top 10 most significant species in the interpolated data are highlighted. Dot colors denote concordance between the abundance coefficients in the original and the interpolated data. **(c, d)** Lead-lag relationships across all species in children with and without ECC. Statistically significant results are plotted in red and blue. Red denotes that a row species significantly leads the column species, and vice versa for blue. Darker colors indicate stronger relationships. **(e)** Hierarchical clustering results after stacking time points and performing ICA.

We perform a time-to-event analysis to identify the hazard for each species abundance to the onset of ECC using a joint model for longitudinal and time-to-event data (4.6.2). We observed a drastic increase in significance after interpolation (Figure 4B), detecting 40 significant species with BGOB (FDR*<* 0.1) compared to just 2 when using the original data. This large microbial community is likely indirectly related through cross-species interactions and lead-lag relationships with the truly causal species and reflects the extensive dysbiosis characterizing the initiation of ECC. Looking at the most significant microbes, we observe several known associations, such as the previously mentioned *Veillonella sp*. and *S. mutans*, alongside *Prevotella salivae, Prevotella melaninogenica*, and *Prevotella pallens* which have been linked to ECC [40, 1, 41]. *Megasphaera micronuciformis* in children with ECC is higher than in children without caries [42]. *Atopobium parvulum* has been found to be predominant in occlusal lesions [43]. Among the previously identified species as differentially abundant, all are significantly associated in the time-to-event analysis, with hazards consistent with the signs of their coefficients in the DA analysis (e.g., *Porphyromonas pasteri* has a negative coefficient in DA and a negative hazard coefficient −59.8). In particular, some novel species such as *Propionibacterium acnes* had the highest hazard coefficients (5394), an order of magnitude higher than some predominantly observed associations (e.g., *S. mutans* with 122). We reason that this may be due to temporal correlations and interactions among truly causal species and that many microbiota may play an indirect role in the onset of the disease, as recently demonstrated with the pathobiont *Selenomonas sputigena* [1].

To interrogate microbe-microbe interactions, we further examine lead-lag relationships across microbes using a signature-based metric (4.6.3). We note the difficulty of this analysis in non-interpolated data, as consistent and numerous intervals are required. We observe 1907 of these relationships between the 92 species that are consistent for at least 70% of individuals, reflecting the broad competition and symbiosis occurring in the oral microbiome. An illustrative example of 16 subjects is shown in Supplementary Figure 1, where *Corynebacterium durum* consistently leads the *S. mutans* group in 101 out of 132 subjects. When comparing patients who developed ECC versus those who did not, we observe a substantial difference (Figures 4C, 4D), with 1844 relationships in non-ECC vs. 2167 in ECC (*P* = 1.97 × 10^−12^). We observe many of the species indicated by previous analyses to have the most drastic change between ECC and non-ECC. The *S. mutans* leads 32 species in the presence of ECC, substantially more than the 14 without, potentially inferring its well-characterized role in ECC could be due to both the species’ own effect and its influence on other species’ dynamics. *Atopobium parvulum* and *Mogibacterium neglectum* lead 9 and 4 species in the presence of ECC, respectively, compared to 0 without. These include several species also previously shown to be associated, including *Stomatobaculum longum, Megasphaera micronuciformis*, and *Actinomyces sp. oral taxon 180*. We also observe *Propionibacterium acnes* leads 18 species in the presence of ECC, and only 4 in non-ECC, many of which were established in previous analyses to have potential relevance, which could explain its disproportionately high hazard. The underlying microbial interaction network may play an important role in the development of ECC and warrants further study.

We further identify temporally correlated groups in the presence of ECC. Interpolation enables stacking of all time points for all individuals into a single matrix that represents how each microbe changes over time. We decompose this matrix into its independent components using ICA and apply hierarchical clustering to identify groups of microbes with similar temporal patterns (Figure 4E). At the lowest resolution, we see a division into two broad clusters. The first, containing 78% of species-level taxa, represents the diversity of the child oral microbiome and contains a range of species mainly associated with health, but also some taxa associated with disease. Bacteria in Cluster 1 include *Corynebacterium durum*, that in conjunction with *Corynebacterium matruchotii* (see below), are now considered keystone bacterial species in the structural development of healthy supragingival plaque. The remaining 22% of species-level taxa form Cluster 2 and include *Actinomyces sp*., *Porphyromonas sp*., *Abiotrophia sp*. and *Neisseria sp*. [44]. Significantly in Cluster 2 we observe two distinct sub-clusters forming: one (sub-cluster 2a) containing species associated with children who did not develop ECC such as *Corynebacterium matruchotii* [45, 40], *Neisseria subflava, Porphyromonas pasteri*, and *Fusobacterium periodonticum* [37], while the other (sub-cluster 2b) comprises species found to have higher abundance in ECC, such as *Veillonella sp*. [25, 26, 27, 28, 29, 30, 31, 32, 46, 47], *Granuticatella elegans, Abitrophia defectiva* and *Lautropia mirabilis* [48, 49]. These sub-clusters appear to be biologically relevant and may reflect the co-varying groups of bacteria that develop in health and disease. Speculatively, the presence, abundance and/or timing of colonization of the keystone species *C. matruchotii* in the health-associated sub-cluster 2a may indicate this bacterium is more relevant for the assembly of a healthy bacterial community than *C. durum* and failure of this species to colonize and proliferate may potentiate susceptibility to ECC.

Thus, BGOB’s interpolation enables a wide variety of well-powered longitudinal analyses, through which we show that ECC onset is likely influenced by the interaction of many microbial species in a dysbiotic microbial community, suggesting a broad set of potential therapeutic targets.

### 2.4 Dynamic Gut Microbe Populations in IBD

We next examine the Inflammatory Bowel Disease Multi’omics Database (IBDMDB, i.e. IBD dataset) [8], which comprises longitudinal gut microbiome data from 130 individuals, 65 with Ulcerative Colitis (UC), 38 with Crohn’s Disease (CD), and 27 healthy patients. The number of time points per individual varies, with 1 to 26 visits each, over a period of up to 57 weeks (Figure 1D). The initial dataset consists of compositional counts and includes 503 microbial species. After quality control, 60 species are retained (4.5.2). We interpolate this dataset using BGOB with a weekly interval, resulting in 58 time points per individual.

We examine differential abundance in the same manner as in the DA analysis of ECC. We observed five significantly associated species (FDR *<* 0.1), compared to 0 in the original data (Figure 5A). We note that the original study was unable to detect any differentially abundant species, even at FDR 0.25 [8], similar to our method when using the original data. The identified species are consistent with known associations. *Parabacteroides* and *Oscillibacter* has been shown to be relevant using similar data [50, 51]. *Eubacterium siraeum* is known to trigger intestinal inflammation [52]. *Subdoligranulum* was further investigated in the original study and found to be of relevance [8]. The *Odoribacter* genus has been found to be negatively associated with IBD [53], although here we find a positive association of *Odoribacter splanchnicus*, highlighting the importance of subgenus granularity and within-genus diversity. We again observe consistency between the signs of the effect sizes of significant species in the original and interpolated data.

**Figure 5.**
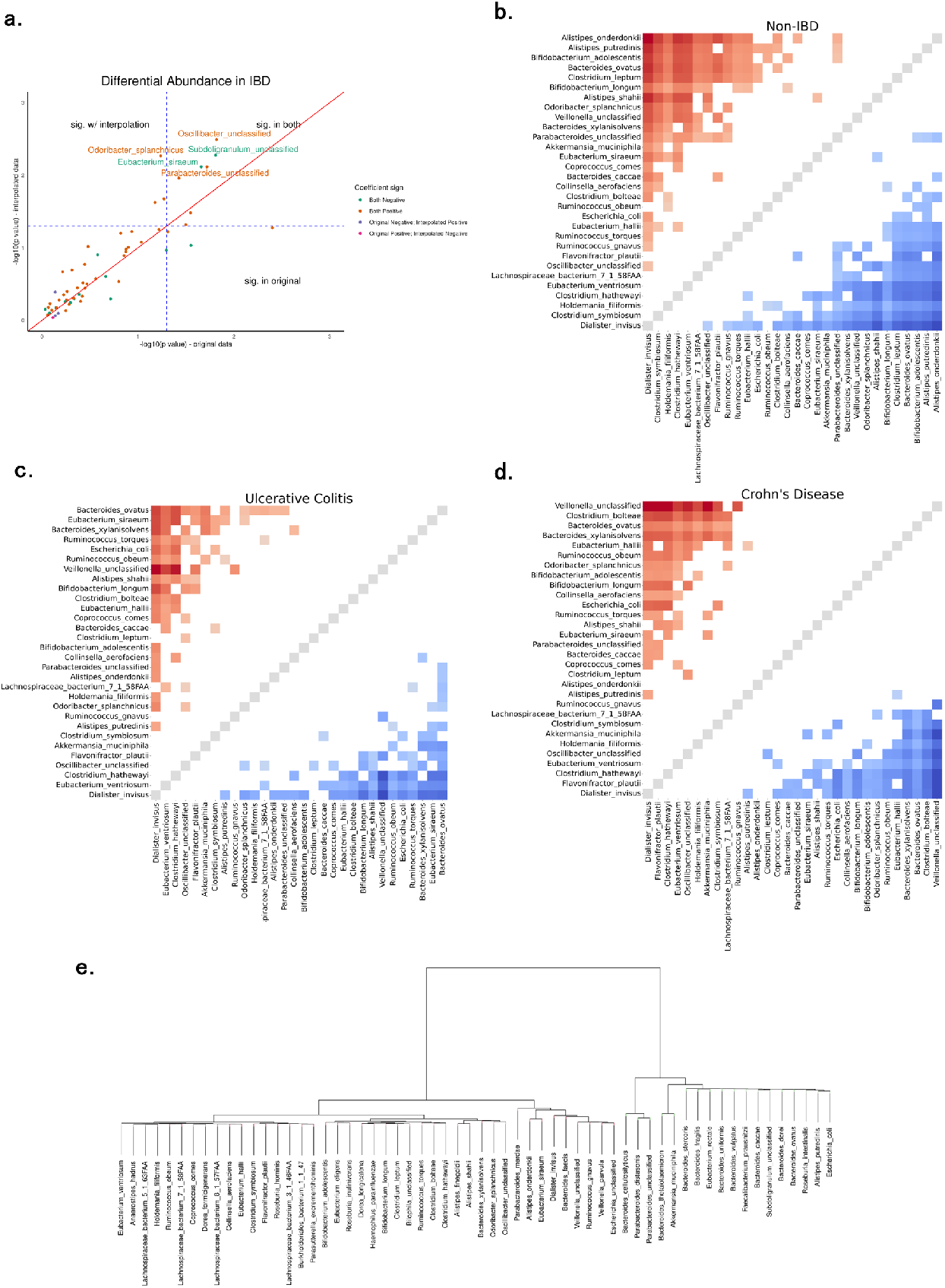
**(a)** Differential abundance results using original data (x-axis) versus BGOB’s interpolated data (y-axis). Species with p-values less than 0.05 on either data are highlighted. Dot colors denote concordance with abundance coefficients in original and interpolated. **(b, c, d)** Lead-lag relationships across all species in patients without IBD, with UC, and with CD. Significant taxa are plotted in red and blue. Red denotes the row species significantly leads the column species, vice versa for blue. Darker colors indicate a stronger relationship. **(e)** Hierarchical clustering results after stacking time points and performing ICA.

Using lead-lag analysis, we observe far less omni-microbial interaction than in ECC (Figures 5B, 5C, 5D). In CD and UC, there are 88 and 79 significant relationships, respectively, out of 1740 possible. Furthermore, in patients without IBD there are 137 relationships, significantly more than in IBD (*P* = 7.3 × 10^−4^ for CD, *P* = 4.6 × 10^−5^ for UC). The species with the largest decreases in how many other microbiota it leads include *Alistipes onderdonkii* (14 healthy, 1 UC, 0 CD) and *Alistipes putredinis* (14 healthy, 1 UC, 1 CD), core microbiota of healthy individuals [54, 55], alongside *Bifidobacterium adolescentis* (14 healthy, 1 UC, 5 CD), which has anti-inflammatory effects and has been developed as a probiotic [56]. This could implicate that several key species lose their microbial regulatory functions in the presence of IBD, even though they themselves may not be differentially abundant. This is further corroborated by the presence of microbes that are leading known pathogenic bacteria and IBD-relevant bacteria *E. coli* in UC and CD, but not in healthy individuals. Disruption in the microbiome regulatory network among healthy and disease-causing microbes could be a potential factor in IBD.

We apply the same temporal clustering to all taxa in the dataset (Figure 5E), resulting in two large clusters. The first cluster, constituting 42 (70%) of species, is potentially pro-inflammatory, due to the presence of microbes such as *Ruminococcus gnavus* [57], *Escherichia coli* [58], and *Clostridium bolteae* [59], which are known to proliferate during inflammatory conditions and produce pro-inflammatory metabolites such as polysaccharides and LPS. Several species in this cluster increase in abundance during active IBD flares and are positively associated with fecal calprotectin levels [60]. The second cluster, comprising 18 (30%) of species, includes anti-inflammatory taxa such as *Faecalibacterium prausnitzii* [61], *Roseburia intestinalis* [62], and *Akkermansia muciniphila*. These species are recognized for their roles in the production of short-chain fatty acids, supporting the integrity of the mucosal barrier, and maintaining immune tolerance. The evaluation of pathway enrichment within cluster 1 and cluster 2 corroborates the pattern of the pro / anti-inflammatory cluster (Supplementary Figure 3). In the first cluster, we observe enrichment of several pro-inflammatory pathways, such as fatty acid degradation [63], phosphotransferase system [64], and degradation of aromatic compounds [65]. In the second, anti-inflammatory pathways are enriched, including vitamin B12 transport and metabolism [66], antibiotic synthesis [67], and lysosomes [68]. This indicates that the temporal correlation of microbial abundances in the gut may be driven by their pro- or anti-inflammation potential, and a temporal microbial network of inflammation regulation may exist.

Overall, interpolation handles the extreme variation of data missingness well and substantially improves power. Downstream analyses reaffirmed known microbiology and identified microbial species with potentially important interactions in the context of IBD.

## 3 Discussion

Previous work on longitudinal microbiome interpolation has focused mainly on statistical methods, such as spline-based approaches, which may overlook the complex interactions among members of the microbial community. Our study demonstrates the efficacy of deep-learning approaches in this context that are capable of modeling continuous time intervals without any distributional assumptions. Accordingly, we introduce BGOB, a novel deep-learning model specifically designed for longitudinal microbiome interpolation, which offers three key advancements over existing methods. First, BGOB flexibly combines interpolation of missing time points (i.e., “unavailable” data) with imputation of zero values in observed data (i.e., “undetectable” data), addressing the challenges that are pervasive in longitudinal microbiome investigations. Second, it incorporates a bidirectional neural-ODE component, enabling accurate predictions across all time points based on user-specified requirements. Third, BGOB is tailored to microbiome data, ensuring biologically meaningful outputs that align with the unique characteristics of microbiome studies. Despite being a deep-learning approach, BGOB does not require extremely large sample sizes and overfitting because the task at hand is reconstruction rather than prediction, and computational requirements are minimal–a CPU is sufficient, and while GPU acceleration is provided, it is not necessary.

Leveraging these advances, we showcase the utility of BGOB through the downstream analyses it enables and improves, including differential abundance testing, time-to-event analysis, lead-lag detection, and clustering. All these analyses are either made possible or greatly enhanced by using BGOB’s interpolated data. Through interpolation, we have overcome power limitations inherent in the original studies and report broad and disease-relevant microbiology. By interpolating oral microbiomes in ECC and gut microbiomes in IBD, both exhibiting various temporal irregularities and count structures, we are able to reveal interacting microbial communities relevant to health and disease, which are biologically informative and have translational potential.

We note several limitations that motivate future work. First, BGOB does not explicitly retain zero-inflated data structures; however, impact on downstream analyses is minimal as interpolated values for true zeros remain small, and the provided package includes post-processing functionality to retain zero-inflation. Second, microbial taxa analyzed here are not exhaustive for the communities under study and will continue to grow as sequencing methods increase in resolution. Third, the analyses performed do not infer causality, but each taxon may be part of a larger, interacting system, which is correlated with truly causal taxa. Looking ahead, the time-to-event framework provided may be a starting point for work that describes causal microbe-disease associations. The lead-lag analysis could be extended to formally incorporate hypothesis testing and to construct microbiome regulatory networks. In the long term, with the increasing success of longitudinal microbiome research, robust interpolation methods will be critical to maximize biological discovery across a diverse array of diseases.

## 4 Methods

### 4.1 Problem Statement

Consider a *N* -subject longitudinal dataset with *V* -dimensional microbial features (i.e., species). We use *j* ∈ {1,2,…,*N*} to denote the *j*-th subject. For the *j*-th subject with *L*_*j*_ time points, we use 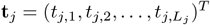 to denote the observed time points. And for *t*_*j,i*_, the *i*-th (*i* ∈ {1, 2, …, *L*_*j*_}) time point in subject *j*, the observation values **y**_**j**,**i**_ will be a *V* -dimensional vector, where the *v*-th feature is denoted as *y*_*j,i,v*_. Therefore, for the *j*-th subject, we can denote the observation values as 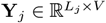.We also introduce **T** = (*T*_1_, *T*_2_, …, *T*_*L*_)^*T*^ to represent all the requisite prediction time points as (1), where *T*_*L*_ is pre-defined and satisfies 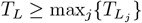.

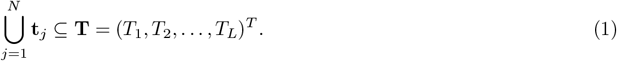

The set of all time points, **T**, includes all possible time points across all subjects 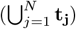.It is important to note, however, that the time points associated with each subject do not necessarily encompass all the time points in **T**. Furthermore, all time points within the set **T** maintain a consistent interval. To more effectively depict the relationship between the *i*-th time point and the *j*-th subject, we introduce a new notation: (*j, i*). This notation denotes the index of the corresponding time points in **T**, such that *T*_(*j,i*)_ = *t*_*j,i*_. Note that a single time point in **T** can be represented in multiple ways. For instance, if *t*_*j,i*_ = *t*_*n,m*_, then *T*_(*j,i*)_ and *T*_(*n,m*)_ are the same.

In this paper, we address two types of missing data to be interpolated: unavailable data and undetectable data. We define unavailable data as instances where no observations are recorded for a subject *j* at a specific time point. For example, for a certain subject, if the specimen is not collected at that time point, there are no corresponding data for all species for this subject. On the other hand, undetectable data refer to cases where the *v*-th observation at a specific time point *t*_*j,i*_ is recorded as zero (i.e, *y*_*j,i,v*_ = 0). This may occur for multiple reasons, such as technical limitations, insufficient sequencing depth, and true zeros. To denote undetectable status for the subject *j*, we utilize a mask vector, **m**_**j**,**i**_ ∈ {0, 1}^*V*^, where a zero entry of *m*_*j,i,v*_ = 0 indicates a lack of data (undetectable data) in the *v*-th dimension at the *i*-th time point for the subject *j*.

### 4.2 Existing Methods

Neural Networks (NNs) are adept at modeling intricate, non-linear relationships between input and output data, making them highly effective in various data analysis tasks. Among these, Recurrent Neural Networks (RNNs) are notably efficient in analyzing longitudinal data due to their inherent design, which captures temporal dependencies. However, a limitation of RNNs is their inherent assumption of evenly spaced time intervals in data sequences. This assumption can hinder their performance on irregularly sampled datasets. To address this, the integration of NNs with ordinary differential equations (ODEs), known as the neural ODE framework, offers a solution. Neural ODEs [20] excel in handling longitudinal datasets characterized by irregular time intervals, providing a continuous-time model that overcomes the discrete-time limitations of traditional RNNs. This approach allows for a more accurate representation of data dynamics in scenarios with irregular temporal sampling.

The GRU-ODE-Bayes (GOB) model [21] is one type of neural ODE model that effectively integrates ODE and GRU. For the *i*-th time point for subject *j*, where the time point is *T*_(*j,i*)_, it employs a modified GRU, which is a type of NN, to model the derivative of the hidden state, *d***h**(*T*_(*j,i*)_). The modified GRU model is called “GRU ODE Cell”, whose implementation details are shown in Supplementary Note. Here, the hidden state **h** represents a vector that captures the temporal dynamics. And for any *T*_(*j,i*)_ + *ϵ*, where 0 *< ϵ* ≤ *T*_(*j,i*+1)_ − *T*_(*j,i*)_, it uses an ODE solver to predict *T*_(*j,i*)_ + *ϵ*, based on *T*_(*j,i*)_, as represented by (2).

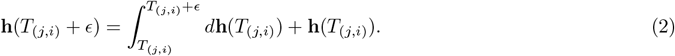

Upon the estimation of **h**(*T*_*i*_) at any *T*_*i*_ by (2), GOB uses multiple layers of NN to predict the values of the observed microbial features **p**(*T*_*i*_) and the corresponding log-variance **lv**(*T*_*i*_) as **p**(*T*_*i*_) = *g*_*p*_(**h**(*T*_*i*_)) and **lv**(*T*_*i*_) = *g*_*lv*_(**h**(*T*_*i*_)), where *g*_*p*_ and *g*_*lv*_ denote the prediction functions, both comprising multiple layers of NN. The implementation details of *g*_*p*_ and *g*_*lv*_ are listed in the Supplementary Note. The observations are fed into the model in time order. For the *j*-th subject at time *T*_*i*_, if the parameters in the hidden state have not been integrated with observation values, we denote it as prior hidden states **h**(*T*_*i*,−_). Subsequently, predicted values and log-variance calculated on the basis of **h**(*T*_*i*,−_) can be expressed as **p**(*T*_*i*,−_) and **lv**(*T*_*i*,−_) respectively. Similarly to Bayesian statistics, we consider this as “prior” information. Once these parameters are integrated with observation values **y**_*j,i*_ using “GRU Observation Cell”, whose implementation details are shown in the Supplementary Note, the prior hidden states turn into “posterior” hidden states **h**(*T*_*i*,+_). Similarly, the predicted values and log-variance on “posterior” hidden states are denoted by **p**(*T*_*i*,+_) and **lv**(*T*_*i*,+_). If the information is unavailable at certain time *T*_*i*_, the values of **h**(*T*_*i*,+_), **p**(*T*_*i*,+_) and **lv**(*T*_*i*,+_) default to **h**(*T*_*i*,−_), **p**(*T*_*i*,−_) and **lv**(*T*_*i*,−_) respectively, which indicates that there is no information for a certain subject at *T*_*i*_.

For any *i* ∈ {1, 2, · · ·, *L*}, the information can be propagated from *T*_*i*−1,+_ to *T*_*i*,+_ using Algorithm 1. Note the subject index *j* is omitted for brevity in the algorithm. Since the algorithm is recursive, for the *j*-th subject, we can initiate **h**(*T*_0,+_) and **p**(*T*_0,+_) and then all the values of the longitudinal data are automatically predicted.

#### Algorithm 1

Information Propagation Algorithm (*j* is omitted for brevity)

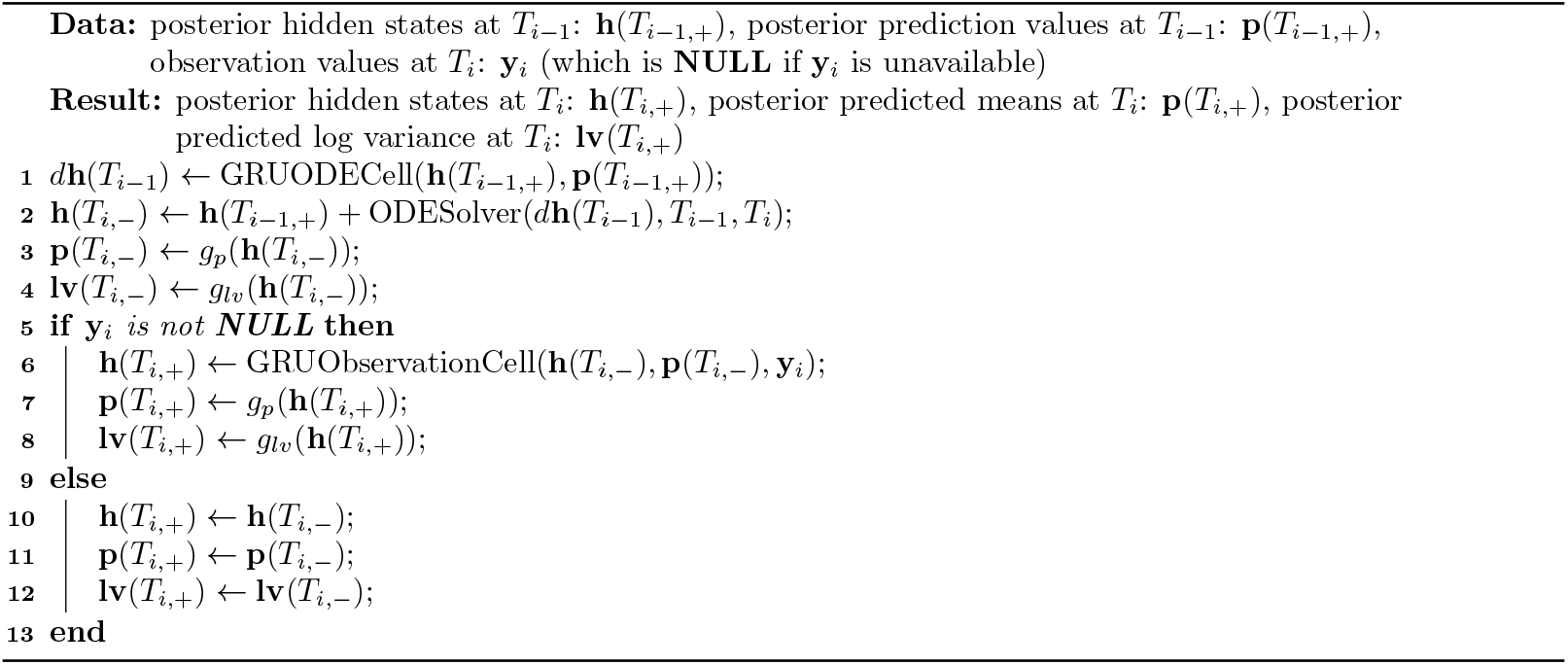

### 4.3 The Proposed Bidirectional GRU-ODE-Bayes Model

The original GOB model faces challenges in producing reliable interpolation results, particularly at early time points, indicating insufficient data for accurate trend estimation. Moreover, the model is not tailored for microbiome data, often resulting in unrealistic outputs, such as negative values. To overcome these limitations, we propose the Bidirectional GRU-ODE-Bayes (BGOB) model, specifically designed for effective interpolation in longitudinal microbiome data.

#### 4.3.1 The Structure of BGOB Model

The model structure of the BGOB model is shown in Figure 2. In this bidirectional model, the information is propagated forward and backward. We use the superscript → and ← to denote them separately. First, we initialize the hidden states at *T*_0_ and *T*_*L*_, denoted as **h**^(→)^(*T*_0,+_) and **h**^(←)^(*T*_*L*,+_), respectively. The hidden states are then calculated and updated by propagating forward and backward through Algorithm 1, resulting in the sequences **h**^(→)^ and **h**^(←)^. By applying the information propagation algorithm in both directions, we obtain the hidden states for all time points in **T**.

For any time points *T*_*i*_ ∈ **T**, in order to calculate predicted values and log-variance at both directions, i.e. **p**^(→)^, **p**^(←)^ and **lv**^(→)^, **lv**^(←)^, we define prediction functions (*g*_*p*_ and *g*_*lv*_) and apply (3) to (6). Specifically designed for microbiome datasets, the prediction functions are structured with five distinct layers, which are listed in Supplementary Note. Note that the final layer in the *g*_*p*_ is a sigmoid function. The inclusion of the sigmoid function in the final layer ensures that the model’s output is constrained between 0 and 1, thereby avoiding the production of nonsensical results.

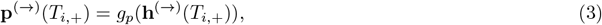

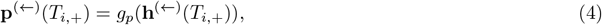

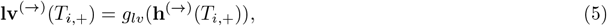

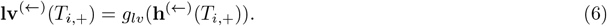

To integrate the information from both forward and backward propagation, at each time point *T*_*i*_ ∈ **T**, we first compute *α*_*i*_ using 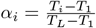, where *α*_*i*_ can be interpreted as a “sureness” parameter. In the forward propagation process, a larger *T*_*i*_ indicates more accumulated information, resulting in greater confidence that the prediction is correct. We then use (7) to obtain the prediction values at time *T*_*i*_, based on the predictions from the forward propagation, **p**^(→)^(*T*_*i*,+_), and the backward propagation, **p**^(←)^(*T*_*i*,+_). The final prediction at *T*_*i*_ can be expressed by (7).

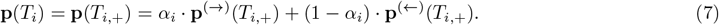

#### 4.3.2 The “Bayesian-like” Loss Function of the BGOB Model

##### The construction of the loss function

The loss function of BGOB consists of two components. The first component, *L*_(*pri*)_, updates the parameters before data integration (prior), while the second component, *L*_(*post*)_, refines the parameters after data integration (posterior).

In order to construct *L*_(*pri*)_, we first define *α*_(*j,i*)_ as 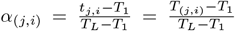, which also represents as the “sureness” of one prediction. *L*_(*pri*)_ in the forward and backward propagation is calculated using (8) and (9). In these equations, ℙ^(*vec*)^(**a**|**b, c**) represents a vector, where the *i*-th element corresponds to the normal density *N* (**a**_*i*_|**b**_*i*_, **c**_*i*_).

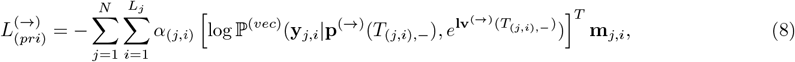

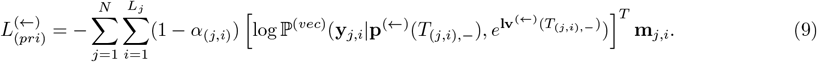

*L*_(*post*)_ represents the error generated after integrating the observed data. To measure the difference between the predicted and the observed data, we use Kullback Leibler (KL) divergence. (10) presents the KL between two normal distributions *N* (*µ*_**1**_, **Σ**_**1**_) and *N* (*µ*_**2**_, **Σ**_**2**_), where *µ*_1_ and *µ*_2_ are the means of the two distributions, and **Σ**_1_ and **Σ**_2_ are their respective covariance matrices, and *n* is the dimension of the random variable.

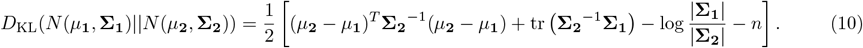

*L*_(*post*)_ in the forward and backward propagation is calculated using (11) and (12), where 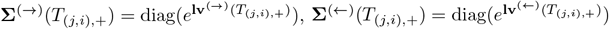, *s* is a pre-defined small constant and **I** is a *V* -dimensional identity matrix.

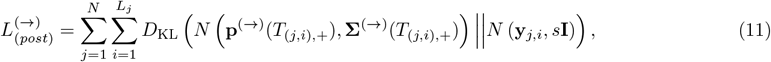

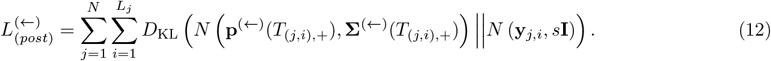

Then we define 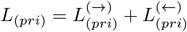, and 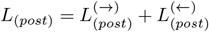. The ultimate loss function combines *L*_(*pri*)_ and *L*_(*post*)_, as shown in (13), where *λ* is a hyperparameter that balances the impact of *L*_(*pri*)_ and *L*_(*post*)_. We utilized the Adam optimizer [69] to minimize the loss *L*. After training the model, we can use (7) to infer the prediction at every time point in **T**.

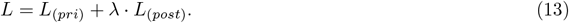

##### Another interpretation of the loss function based on “ELBO”

In addition to its Bayesian-inspired interpretation, the BGOB loss function can also be understood through the lens of the Evidence Lower BOund (ELBO), as constructed in the Variational Autoencoder (VAE) loss function [70]. Consider the *i*-th observation of subject *j*: if **y**_*j,i*_ is available, we denote the true distribution of the *V* microbiome species as **Y**_*j,i*_, the distribution of the hidden state before integrating the actual observation as **H**_*j,i*,−_, and the updated hidden state after integrating the observed data as **H**_*j,i*,+_. For simplicity, we omit the subscripts *j, i* in this section.

In the original VAE model, the goal is to map one **Y** to another distribution **Z** and then back to the original space. The loss function, i.e. ELBO, comprises a reconstruction error and a prior matching error. Inspired by this framework, we design similar loss functions tailored for our model. We assume that the hidden state **H**, in both forward and backward propagation, captures the temporal information preceding each time point. Therefore, our loss function is constructed with two main objectives: First, the prior hidden state **H**_−_ should accurately predict the observed microbiome data **Y**. Additionally, the integration of the observed data **Y** should effectively update the hidden state to **H**_+_, reflecting the additional information gained from the observations.

Therefore, similar to the VAE model, we assume that the encoder distribution **Y**|**H**_−_ follows a normal istribution *N* (*µ*_*ϕ*_(**h**), **Σ**_*ϕ*_(**h**)). We use the negative log-likelihood to construct the encoder loss. Specifically, in the forward propagation, *µ*_*ϕ*_(**h**) corresponds to **p**^(→)^(*T*_(*j,i*),−_), and in the backward propagation, it corresponds to **p**^(←)^(*T*_(*j,i*),−_). Similarly, the covariance matrix **Σ**_*ϕ*_(**h**) is represented by 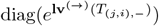 during forward propagation and by 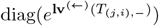 during backward propagation. Incorporating the “sureness” parameter *α*, and recognizing that undetectable species do not contribute to the final loss, we clearly express the resulting equations for both forward and backward propagation as shown in (8) and (9). Adding up (8) and (9), we get *L*_(*pri*)_, which is like the reconstruction error in VAE.

Additionally, we assume that the decoder distribution **H**_+_|**Y** follows a normal distribution *N* (*µ*_*θ*_(**y**), **Σ**_*θ*_(**y**)) and use the Kullback-Leibler (KL) divergence to construct the decoder loss. In this context, *µ*_*θ*_(**y**) corresponds to **p**^(→)^(*T*_(*j,i*),+_) during forward propagation and **p**^(←)^(*T*_(*j,i*),+_) during backward propagation. The covariance **Σ**_*θ*_(**y**) is given by 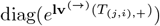 in the forward direction and by 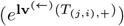 in the backward direction. Taking into account the lack of contribution from undetectable species, we derive explicit equations for both forward and backward propagation, as presented in (11) and (12). Adding up (11) and (12), we get *L*_(*post*)_, which is like the prior matching error in VAE. We also introduce a hyperparameter *λ* to better balance the training process of *θ* and *ϕ*, resulting in the final loss function in (13).

Thus, despite the apparent complexity of the final loss function, it is fully interpretable. The first component, *L*_(*pri*)_, functions similarly to a reconstruction error, tuning the model parameters before the integration of real observations. The second component, *L*_(*post*)_, acts as a prior matching error, refining the parameters after the integration of observed data. Together, *L*_(*pri*)_ and *L*_(*post*)_ create an loss function similar to ELBO, effectively balancing the learning of prior knowledge with adjustments driven by actual data.

### 4.4 Simulation Procedures

#### 4.4.1 Univariate Simulation Procedures

We use zero-inflated log-normal (ZILN) distribution to simulate microbiome data. First, we define a log-normal distribution with mean *µ* and variance *µ*^2^*θ*^2^, which is denoted as *LN* (*µ, µ*^2^*θ*^2^). If *X* ∼ *LN* (*µ, µ*^2^*θ*^2^), its probability density function can be written as (14), 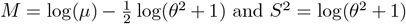. And (15) denotes the probability function of a ZILN distribution.

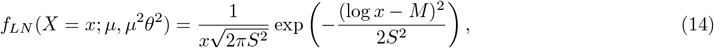

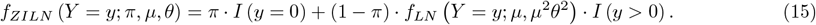

Since the ZILN model can only generate data at one time point, we need to simulate the mean for subject *j* at time *t*, i.e. *µ*_*j,t*_, to generate different longitudinal trends. We introduce two stochastic processes to generate different *µ*_*j,t*_. We first define 1-dimensional Brownian motion *W*_*t*_, whose differential *dW*_*t*_ is defined by *dW*_*t*_ ∼ *N* (0, *dt*), where *N* (*µ, σ*^2^) denotes normal distribution with mean *µ* and standard deviation *σ*, and *dt* denotes the derivative of time *t*. And we employ the Black-Scholes and Ornstein-Uhlenbeck processes to generate varying trends for *µ*_*j,t*_. The corresponding stochastic differential equations for these processes are presented in (16) and 17, where *k, σ*, and *m* are tuning parameters.

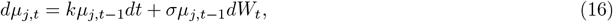

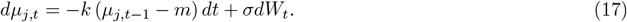

To simulate longitudinal datasets with different trends, we define *P* distinct groups, each following a unique trend. These trends are generated using varied models and diverse parameter settings for *k, σ*, and *m*. For the *j*-th subject at time *t*, the means of these trends are denoted as 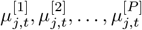. The approach to generate microbiome longitudinal data with a diverse range of temporal patterns as follows: For each subject *j*, we assign them to a group *p* ∈ 1, 2, …, *P*. And for *t* ∈ {1, 2, · · ·, *T*}, we first generate 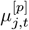 based on the group membership *p* and then simulate *y*_*j,t*_ to follow a ZILN distribution characterized by parameters 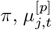, and *θ*. The zeroes generated during this step represent undetectable data. Additionally, data points are randomly dropped at each time points with a probability of *δ*, thus creating unavailable data. Therefore, after the procedure of introducing missing data, the proportions of unavailable data, undetectable data, and remaining information are approximately *π* · (1 − *δ*), *δ*, and (1 − *π*) · (1 − *δ*) (denoted as *η*), respectively.

In our simulation setup, we define 5 distinct groups (*P* = 5), each representing a unique time trend for *µ*_*j,t*_, with 200 subjects per group. For each subject, we simulate *T* = 30 time points. The specific models and parameters used to generate *µ*_*j,t*_ for each group are provided in the Supplementary Note. Additionally, we set 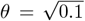. To evaluate the robustness of our method under varying levels of missing data, we vary *π* and *δ* across different scenarios. Here, *π* governs the proportion of undetectable data, while *δ* controls the proportion of unavailable data. The settings for *π* and *δ* in the different scenarios are listed in the Supplementary Note.

#### 4.4.2 Multivariate Simulation Procedures

In the multivariate longitudinal data simulation, in order to simulate both the intra- and inter-microbiome interactions, we use a method inspired by generalized Lotka–Volterra (gLV) model [24]. Specifically, as shown in Figure 6A, for a single time point *t* for subject *j*, we can separate the increment of microbiome *v* (*dµ*_*j,t,v*_) as two resources, which are 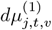 and 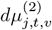, representing the change rate caused by intra-microbiome interaction and inter-microbiome interaction separately.

**Figure 6.**
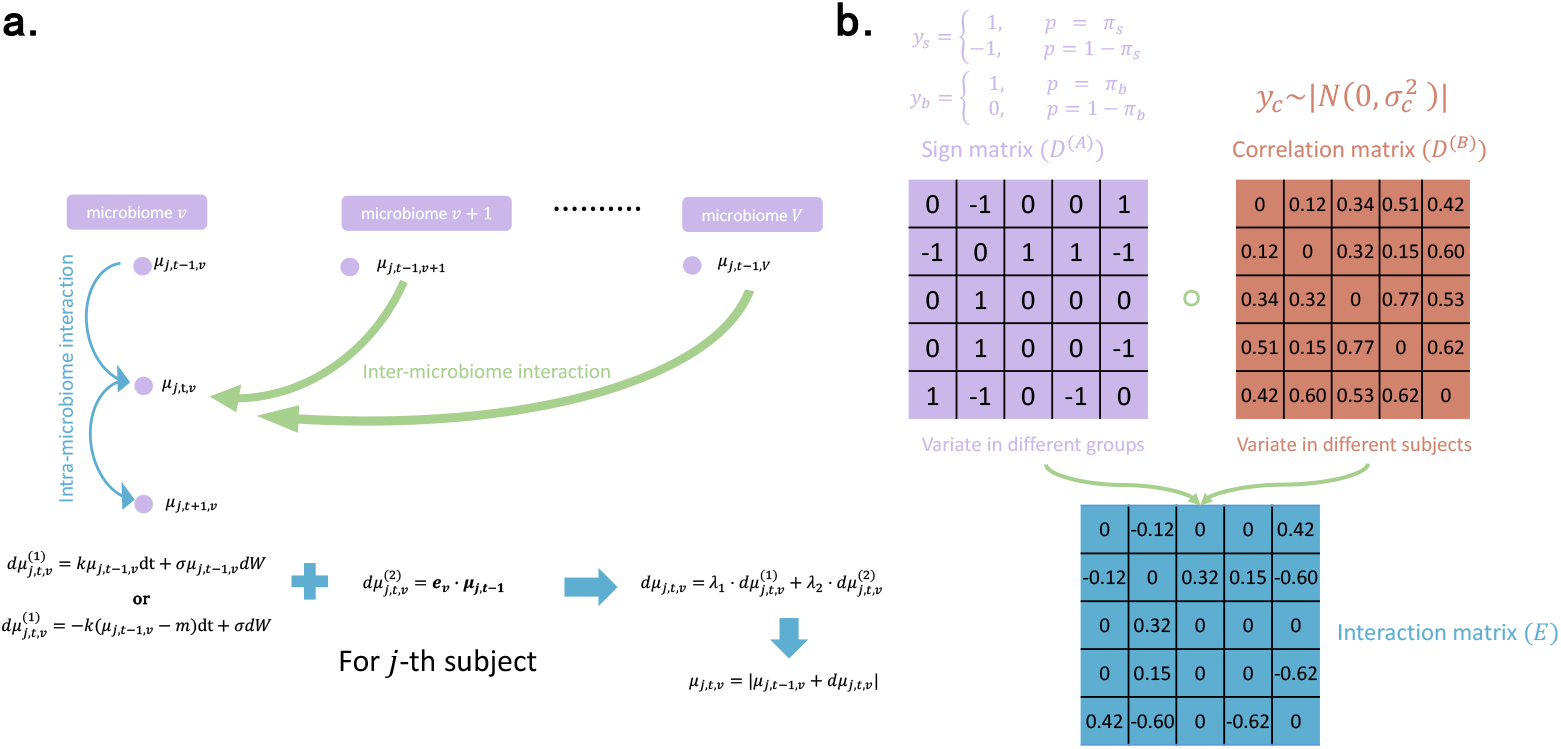
Multivariate simulation setting. **(a)** The interaction is separated as intra- and inter-microbiome interaction. The intra-microbiome interaction is modeled using stochastic process while inter-microbiome interaction is defined by matrix operations. **(b)** The sign matrix determines the direction of the interaction, which is consistent within one group. The correlation matrix determines the extent of the influence, which is consistent within one subject. Therefore, the interaction matrix guarantees that within the same group, the direction of influence stays the same, while the extent of influence may variate for different subjects. And within the same subject, the extent of influence will also stays the same at different time points.

For the intra-microbiome interaction 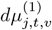, similar to univariate data simulation, we use Black-Scholes process or Ornstein-Uhlenbeck process to model the interaction as (18) and (19) shows, where different microbiomes are simulated using different models or the same model with different parameters in the same group. For the same microbiome in different groups, we still use different models or the same model with different parameters *k, σ* and *m*. This approach generates multiple groups with distinct time trends, effectively representing the interaction by the microbiome itself.

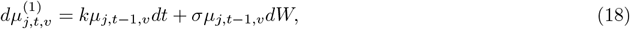

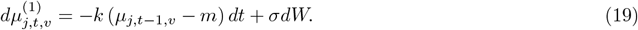

In order to generate inter-microbiome simulation data, we use the method shown in Figure 6B to generate the interaction matrix **E**. Specifically, we begin by creating two matrices: a sign matrix and a correlation matrix. The interaction matrix is then generated using the Hadamard product of the sign matrix and the correlation matrix. The sign matrix **D**^(*A*)^ can be generated through the following three steps. First, we construct the matrix **D**^(*A*)^, where each entry is defined as 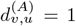 for *v < u*, and 0 otherwise. Next, for every 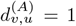 (where *v < u*), we independently modify its value by multiplying it by 1 with probability *π*_*b*_, and by 0 otherwise. Similarly, for each remaining 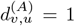 (where *v < u*), we further adjust its value by multiplying it by 1 with probability *π*_s_, and by −1 otherwise. Finally, we symmetrize the matrix and get **D**^(*A*)^. For the correlation matrix **D**^(*B*)^, we fill 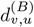 (where *v < u*) with values generated by *y*_*c*_, where 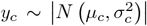. Here, |*N* (*µ*, σ)^2^| represents a Half-Normal distribution derived from a normal distribution with mean *µ* and variance *σ*^2^. The diagonal elements of **D**^(*B*)^ (denoted as 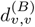) are set to 0. Again, we symmetrize the matrix and get **D**^(*B*)^. After generating matrices **D**^(*A*)^ and **D**^(*B*)^, we generate the interaction matrix **E** using **E** = **D**^(*A*)^ ∘ **D**^(*B*)^. And we calculate 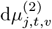 using 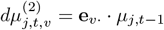, where **e**_*v*·_ denotes the *v*-th row of matrix **E** and *µ*_*j,t*−1_ = (*µ*_*j,t*−1,1_, *µ*_*j,t*−1,2_, …, *µ*_*j,t*−1,*V*_)^*T*^.

Using the aforementioned generation process, the sign matrix can vary across different groups. However, within the same group, the sign matrix remains consistent for all subjects, while the correlation matrix may differ between subjects. This ensures that the direction of the correlation (i.e., the sign) between two microbiome species is preserved within a group, although the magnitude of the correlation may vary slightly between subjects. In contrast, both the magnitude and the sign of the correlation parameters can vary across different groups, allowing for distinct interaction patterns between groups. This approach allows for significant flexibility in the simulation while ensuring that subjects within the same group do not behave too differently. After generating 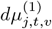 and 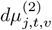 using the proposed methods, we obtain *dµ*_*j,t,v*_ using 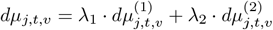, where *λ*_1_ and *λ*_2_ are tuning parameters. Finally, we calculate *µ*_*j,t,v*_ using *µ*_*j,t,v*_ = |*µ*_*j,t*−1,*v*_ + *dµ*_*j,t,v*_|.

To generate the observed data, we assign each subject *j* to a group *p*, where *p* ∈ {1, 2, …, *P*}. For each microbiome *v* with group membership *p*, we generate 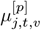 with varying time trends across different groups based on both the intra- and inter-microbiome interaction, and simulate *y*_*j,t,v*_ using a zero-inflated log-normal distribution with parameters 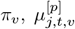, and *θ*. Here, *π*_*v*_ represents the zero-inflation rate for microbiome species *v*, calculated as logit(*π*_*v*_) = *α* · *S*_*v*_ + *β*, where 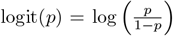, *S*_*v*_ denotes the relative mean abundance of species *v*, and *α* and *β* are hyperparameters. The zeroes generated by this process correspond to undetectable data. Additionally, data is randomly omitted at certain time points with a probability *δ*, introducing unavailable data. This simulation process ensures that if one time point is unavailable for a given subject, all microbiome information for that time point is dropped. However, at some available time points, it’s possible that the information for certain microbiome is undetectable, while the rest of the information for that time point remains. This perfectly emulates the circumstances encountered in real datasets.

In our experiment, we have 5 groups (*P* = 5), each group includes 80 subjects, where each subject has 5 microbiome species. We simulate 30 time points for each subject. The trends for the same microbiome vary among different groups. To generate the inter-microbiome interaction, we first generate matrix *D*^(*A*)^ with *π*_*s*_ = 0.7 and *π*_*b*_ = 0.4. We randomly generate 5 different *D*^(*A*)^ for different groups. For each subject in a specific group, we generate *D*^(*B*)^ with *µ*_*c*_ = 0.1 and *σ*_*c*_ = 0.001. To generate the inter-microbiome interaction, we specify 5 different stochastic processes for each group. And we set 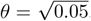. The values of *π*_*v*_ vary among different simulation data sets, as *β* is different between different simulations. The zeroes generated in this step will be undetectable data. We then randomly drop data at certain time points with a probability of *δ*, thus generating unavailable data. We have developed four simulation scenarios, and for each scenario, the parameters *α, β*, and *δ* have been chosen to manage missing information throughout the data sets. Specifically, *α* and *β* are used to adjust the amount of data that is unavailable, while *δ* regulates the proportion of data that is undetectable. The details of these scenarios are shown in the Supplementary Note.

#### 4.4.3 Simulation Evaluation

We created both univariate and multivariate simulation datasets following the procedure described above. For each of these simulations—univariate and multivariate—we first process with a log transformation and then interpolate with three different methods: cubic spline interpolation, the original GOB method, and our BGOB method. And we repeat the simulation 20 times using different random seeds, and evaluate our results based on the mean of these simulations.

We begin by evaluating the accuracy of interpolation in these datasets. This is done using the root mean square error (RMSE) as a measure of prediction accuracy, as detailed in (20).

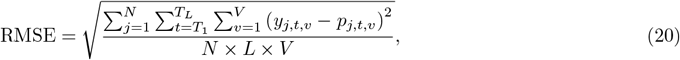

where *y*_*j,t,v*_ denotes the value of *v*-th microbiome of subject *j* at the time point *t* and *p*_*j,t,v*_ denotes the prediction values of the same microbiome of the same subject at the same point. Since we interpolate the data after log transformation, *y*_*j,t,v*_ and *p*_*j,t,v*_ are all in log scale. *N* equals to the number of subjects, *L* equals to the number of total time points after interpolation, and *V* equals to the number of different species. For univariate simulation, we use count data to calculate RMSE since there is no point to calculate the percentage data based on one species. RMSE is calculated based on the percentage data for multivariate simulation.

Next, to assess the clustering efficiency after interpolation, we utilize “Time Series K-Means” clustering in the Python package “tslearn” [71] to cluster the interpolated data using the three different methods. The performance of this clustering was then evaluated by the purity score defined as 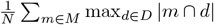, where *D* is the real set of classes and *M* denotes the predicted set of classes using a specific model. Within each simulation, we carry out the clustering procedure using 100 different seeds and calculate the mean purity score to represent the purity score for that simulation.

Additionally, we compare the interpolation accuracy in the multivariate simulation datasets between our BGOB method and two other methods: TGP-CODA [15] and Luminate [16]. Both TGP-CODA and Luminate are Bayesian models specialized in handling multivariate microbiome longitudinal datasets. TGP-CODA utilizes a Gaussian process model tailored to longitudinal count data, considering the temporal correlation of adjacent time points. Additionally, it incorporates a hierarchical model to address overdispersion and introduces a separate parameter for technical zeros. Luminate uses a state-space model to determine relative abundance from longitudinal microbiome data and distinguishes between biological and technical zeros. We still use RMSE to evaluate performance. However, since Luminate does not interpolate missing data, the RMSE is calculated for Luminate exclusively at time points where information is available. Also, since we directly apply TGP-CODA and Luminate to the data without the log-transformation, we transform back our BGOB-interpolated dataset and then compare it with the datasets processed by TGP-CODA and Luminate. Therefore, *y*_*j,t,v*_ and *p*_*j,t,v*_ in (20) are in the original scale instead of the log scale.

### 4.5 Real Data Description

#### 4.5.1 ECC

In this study, we used VicGen early childhood caries (ECC) data [7], a longitudinal cohort study of infants established in 2008 to track their natural history of dental disease and investigate causative factors. The data includes 132 individual child participants with clinical and sequencing data, among whom 42 had ECC (i.e., the clinical endpoint of interest) before their last visit. ECC is a person-level diagnosis defined as one or more dental cavities in a child under the age of 6 [72]. In this study, ECC (i.e., dental cavities) was defined as a caries lesion of ICDAS (International Caries Detection and Assessment System) code 3 or greater [73], consistent with previous investigations [74]. Each participant had multiple visits at around six key time points: the first being unknown and set at 0 months, with the subsequent five averaging at 7.7, 13.2, 19.7, 39.0, and 48.6 months of age. Figure 1C depicts the data collection time points for each individual. In each sample, there is information on 857 species. To ensure biological relevance, we grouped species based on 16S rRNA sequencing and phylogenetic relationships as described by Whiley and Beighton [75]; details of these groupings are provided in the Supplementary Note. Of note, the data are compositional. After quality control, taxa with *>* 80% zeros were excluded, leaving 92 species (including ‘Other’) for analyses.

We also have time-to-event data that largely overlap with the sequencing data. This dataset includes 119 common participants with observations recorded at four distinct time points, averaging 19.7, 39.0, 48.5, and 60.4 months of age. At each of these time points, binary variables indicating whether the event of interest (ECC) has occurred are assessed. In this study, 55 out of 119 participants developed ECC before their final visit. We note the time-to-event data cover a slightly longer period than the sequencing data (60.4 vs 48.6 months, respectively).

#### 4.5.2 IBDMDB

We further demonstrate our method using the Inflammatory Bowel Disease Multi’omics Database (IBDMDB) [8], referred to here as the IBD dataset. This dataset contains information on 130 individuals, 27 of whom do not have Inflammatory Bowel Disease (IBD). Among those diagnosed, 65 have Ulcerative Colitis (UC), and 38 are affected by Crohn’s Disease (CD).

We analyze the Metagenomics (MGX) dataset from this resource. Taxonomic profiles were generated using MetaPhlAn, while functional profiling was performed with HUMAnN2 to quantify gene presence and abundance on a per-species basis (UniRef90s) for both metagenomics and metatranscriptomics. To ensure adequate read depth, samples with at least 1 million reads (after human filtering) were included and at least one non-zero microbial abundance detected by MetaPhlAn. At the species level, the dataset initially contained data for 503 species. The data are compositional, with abundances summing to 1 across all species within each sample. After quality control, which excluded taxa with *>* 70% zeros, 60 species remained for analysis. The frequency of visits per individual varied widely, ranging from 1 to 26 visits, with the longest observed duration extending up to 57 weeks. Additionally, the time intervals between visits were inconsistent. An overview of the time points for these individuals appears in Figure 1D.

### 4.6 Downstream Analysis for Real Data

#### 4.6.1 Differential Abundance (DA) analysis

Differential abundance (DA) analysis quantifies shifts in the composition of the microbial community under different conditions. We perform DA analysis using the “gamm4” package [76] to identify species most significantly associated with ECC or IBD. The gamm4 model is formulated as shown in (21), where *y*_*j,i*_ represents the interpolated abundance of a species using BGOB for the *i*-th time point in the *j*-th subject. *x*_*j*_ is a binary variable indicating the presence or absence of ECC or IBD in the subject, and *t*_*j,i*_ represents time. The term *b*_*j*_ accounts for the random effect of the *j*-th subject, and *β* denotes the regression parameter. The time variable is smoothened with a thin plate regression spline [77], represented by *s*(*t*_*j,i*_).

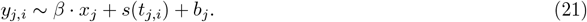

#### 4.6.2 Time-to-Event Analysis

We use a joint model for longitudinal and time-to-event data, where the longitudinal component is inferred using our BGOB model. The Time-Dependent Cox Model [78] is an extension of the standard Cox Proportional Hazards Model [79], allowing covariates to evolve over time. In the hazard function (22), *h*(*t*) represents the hazard at time *t*, and *y*(*t*) denotes the interpolated DNA-based species taxonomic abundance at time *t* for a specific species. The baseline hazard function *h*_0_(*t*) denotes the inherent risk present across all subjects, independent of covariate values. *β* is a set of coefficients to be estimated, each corresponding to one of the species in the study. These coefficients quantify the effect size of the species abundance on the hazard rate, such that a positive *β* reflects a species abundance leading to increased disease risk, and vice versa. We conduct the time-to-event analysis exclusively for the ECC dataset, as the IBD dataset lacks corresponding time-to-event outcomes.

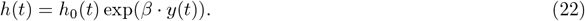

#### 4.6.3 Pairwise Lead–Lag Detection and Network Analysis

A lead-lag relationship is established when the earlier values of a longitudinal trend are more predictive of the later values of another than the reverse [80]. To detect this, we opt for the rough path theory [81], which examines iterated integrals of longitudinal data. Borrowing notation from [82], for a given continuous longitudinal dataset over a time interval [*a, b*], where *a* and *b* represent the start and end times respectively, we consider a path *X* : [*a, b*] → ℝ^*d*^. We denote the coordinate paths of *X* by 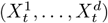 for *t* ∈ [*a, b*], where each 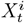 represents the *i*-th component of *X* at time *t*. In our work, a component corresponds to a microbial feature (taxon, species). Thus, each *X*^*i*^ : [*a, b*] → ℝ is a real-valued longitudinal series corresponding to the *i*-th species. The signature *S*(*X*)_*a,b*_ is an infinite series comprising all iterated integrals of *X*. For a multi-index (*i*_1_, …, *i*_*k*_) with *i*_*j*_ ∈ 1, …, *d*, where *d* is the number of species, the superscript 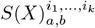 denotes the iterated integral of *X* along the components *i*_1_, …, *i*_*k*_, defined in [82] as (23).

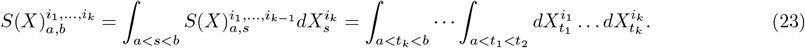

The first level with single index *i* ∈ {1, …, *d*} is 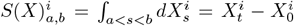, representing the time series increments. The second level with a pair of indices 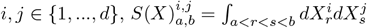 captures the area enclosed by the two series and a straight line connecting their endpoints (Lévy area). To investigate the pairwise lead-lag of two species, we use the Lévy area as the lead-lag metric. The Lévy area is given by 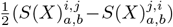, where a positive value indicates that movements in the first component (*X*^*i*^) tend to precede and predict movements in the second (*X*^*j*^) (See [82] for a more complex rationale for the lead-lag metric). We compute these pairwise measures with python library “iisignature” [83].

#### 4.6.4 ICA

Independent component analysis (ICA) is performed on the longitudinal data using the R package “fastICA” [84]. This process transforms raw longitudinal data into sets of independent variables for later cluster analysis to identify sets of features that have similar temporal patterns. Specifically, it uses the independent factors (latent variables) in multivariate data and decomposes the input matrix into statistically independent components. ICA assumes that the observed data **X** = (*x*_1_, *x*_2_, …, *x*_*n*_)^*T*^ within the space ℝ^*n×m*^ can be modeled as a linear combination of *I* independent components **S** = (*s*_1_, *s*_2_, …, *s*_*I*_)^*T*^, with some unknown mixing matrix **A**, i.e., **X**_*n×m*_ = **A**_*n×I*_ · **S**_*I×m*_. Consider the interpolated data set **Y** ∈ ℝ^*N×L×V*^, where *y*_*j,i,v*_ corresponds to the relative abundance of species *v* at time *i* for subject *j*. Since performing ICA separately for each subject can lead to variability in source separation across subjects [85], a common approach, particularly in fMRI analysis, is group ICA [86]. Data from multiple subjects are temporally concatenated, and ICA is then applied to the combined data matrix. To be specific, we first stack each time point in each individual to generate **Y**_(*N*·*L*)*×V*_. We set the number of independent components *I* = 15. The **Y** matrix is then decomposed into group ICA matrix **S**_*I×V*_, where each independent component consists of *V* different species. The flowchart of group ICA is illustrated in Figure 7.

**Figure 7.**
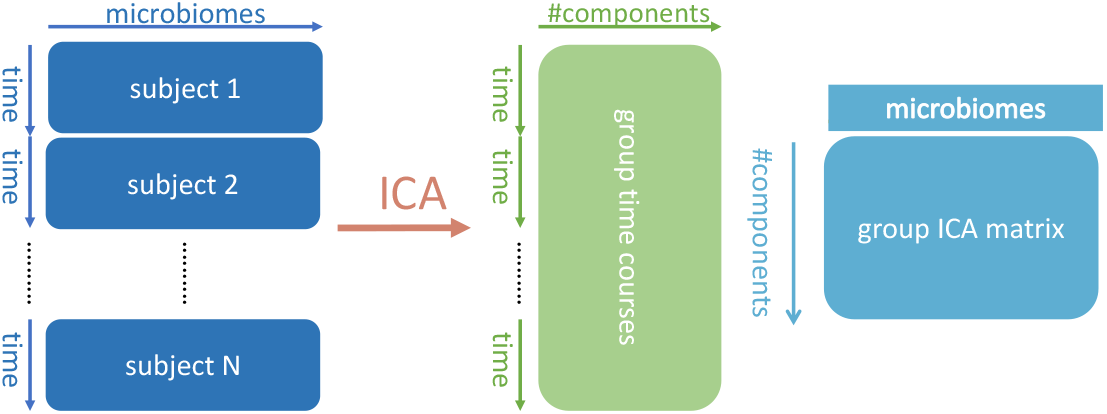
Flowchart of group ICA. All time points of a study participant are stacked consecutively for all microbial taxa, and all participants are then stacked. ICA decomposition is then performed on the entire stacked matrix, giving an independent component by microbiome matrix.

#### 4.6.5 Hierarchical Clustering

We then input **S** into hierarchical cluster analysis using Ward’s method [87]. We first define the increment of two groups A and B as 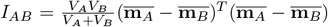, where *V*_*A*_ and *V*_*B*_ represent the number of species in groups A and B, and 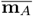 and 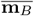 are the mean vectors of the independent components of A and B, calculated from **S**. Specifically, 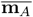 and 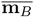 are obtained by averaging the independent components in **S** corresponding to the species grouped in A and B. At each step, we minimize the increment in the within-group error sum of squares by merging the two groups that result in the smallest increase, continuing iteratively until the clustering is completed.

#### 4.6.6 Pathway Enrichment

To inform the clustering results, we perform pathway enrichment for a set of microbial taxa, against all taxa in the dataset. First, we filter out any species not present in KEGG (https://rest.kegg.jp/list/organism). We either match taxa directly by name, or if only higher-level information is available (e.g., *E. coli* versus *E. coli* ‘s strains), we consider all species’ pathways. Having collected all the pathways for all available species, we carry out a Fisher’s Exact Test to determine which pathways are significantly enriched in a subset of species.

