## Supplementary Note for "Deep-learning-based interpolation of longitudinal microbiome data powers biologically informative discovery"

### Appendix for "BGOB: A novel interpolation model for irregularly-sampled microbiome data based on ODE-related deep learning methods"

Yixiang Qu[†1], Ruiqi Lyu[†2], Duan Wang[1,3], Yifan Dai[1], Alistair Turcan[2], Shilin Yu[1], Jialiu Xie[1], Jeffrey Roach[4], Catherine Butler[5], Pew-Thian Yap[6], Hongtu Zhu[1], Stuart Dashper[5], Apoena Aguiar Ribeiro[7], Didong Li[1], Kimon Divaris[8,9], Di Wu[1,10,*]

**1 Department of Biostatistics, Gillings School of Global Public Health, University of North Carolina at Chapel Hill**
**2 School of Computer Science, Carnegie Mellon University**
**3 Chapel Hill High School**
**4 Research Computing, University of North Carolina at Chapel Hill**
**5 Melbourne Dental School, The University of Melbourne**
**6 Department of Radiology, School of Medicine, University of North Carolina at Chapel Hill**
**7 Department of Diagnostic Sciences, Adams School of Dentistry, University of North Carolina at Chapel Hill**
**8 Department of Pediatric Dentistry and Dental Public Health, Adams School of Dentistry, University of North Carolina at Chapel Hill**
**9 Department of Epidemiology, Gillings School of Global Public Health, University of North Carolina at Chapel Hill**
**10 Department of Biomedical Sciences, Adams School of Dentistry, University of North Carolina at Chapel Hill**

*****

### 1 Structure of "GRU ODE Cell"

The "GRU ODE Cell" takes $\mathbf{p}(T_{(j,i),+})$ and $\mathbf{h}(T_{(j,i),+})$ as input and outputs $d\mathbf{h}(T_{(j,i)})$. The entire operation from Eq. 1 to Eq. 4 is represented as $\text{GRUODECell}(\mathbf{h}(T_{(j,i),+}), \mathbf{p}(T_{(j,i),+}))$, where $\sigma$ denotes the sigmoid activation function ($\sigma(x) = \frac{1}{1+e^{-x}}$) and $\mathbf{W}$, $\mathbf{U}$, $\mathbf{b}$ denote parameters.

$$\mathbf{r}(T_{(j,i)+1}) = \sigma\left(\mathbf{W}_r\mathbf{p}(T_{(j,i),+}) + \mathbf{U}_r\mathbf{h}(T_{(j,i),+}) + \mathbf{b}_r\right), \tag{1}$$

$$\mathbf{z}(T_{(j,i)+1}) = \sigma\left(\mathbf{W}_z\mathbf{p}(T_{(j,i),+}) + \mathbf{U}_z\mathbf{h}(T_{(j,i),+}) + \mathbf{b}_z\right), \tag{2}$$

$$\mathbf{g}(T_{(j,i)+1}) = \tanh\left(\mathbf{W}_h\mathbf{p}(T_{(j,i),+}) + \mathbf{U}_h\left(\mathbf{r}(T_{(j,i)+1}) \circ \mathbf{h}(T_{(j,i),+}) + \mathbf{b}_h\right)\right), \tag{3}$$

$$d\mathbf{h}(T_{(j,i)}) = (1 - \mathbf{z}(T_{(j,i)+1})) \circ (\mathbf{g}(T_{(j,i)+1}) - \mathbf{h}(T_{(j,i),+})). \tag{4}$$

### 2 Structure of "GRU Observation Cell"

The Algorithm 1 indicates how the real data $\mathbf{y}_{j,i}$ is integrated into the hidden state. The whole algorithm is denoted as $\text{GRUObservationCell}(\mathbf{h}(T_{(j,i),-}), \mathbf{p}(T_{(j,i),-}), \mathbf{y}_{j,i})$. It takes $\mathbf{h}(T_{(j,i),-})$, $\mathbf{p}(T_{(j,i),-})$, $\mathbf{p}(T_{(j,i),-})$, and $\mathbf{y}_{j,i}$) as input and outputs $\mathbf{h}(T_{(j,i),+})$, $\mathbf{p}(T_{(j,i),+})$, $\mathbf{lv}(T_{(j,i),+})$.

**Algorithm 1:** GRU Observation Cell ($j$ is omitted for brevity)

**Data:** prior hidden states at $T_i$: $\mathbf{h}(T_{i,-})$, prior predicted means at $T_i$: $\mathbf{p}(T_{i,-})$, prior predicted log variance at $T_i$: $\mathbf{lv}(T_{i,-})$, observation values at $T_i$: $\mathbf{y}_i$

**Result:** posterior hidden states at $T_i$: $\mathbf{h}(T_{i,+})$, posterior predicted means at $T_i$: $\mathbf{p}(T_{i,+})$, posterior predicted log variance at $T_i$: $\mathbf{lv}(T_{i,+})$

$\mathbf{er}(T_i) \leftarrow (\mathbf{y}_i - \mathbf{p}(T_{i,-})) \oslash \sqrt{e^{\mathbf{lv}(T_{i,-})}}$;

▷ Here $\mathbf{y}_i$, $\mathbf{p}(T_{i,-})$, $\mathbf{lv}(T_{i,-})$, $\mathbf{er}(T_i)$ are all column vectors, and $concat\,[\mathbf{a}, \mathbf{b}]$ denotes the stack of column vectors with the same size

$\mathbf{info}(T_i) \leftarrow concat\,[\mathbf{y}_i, \mathbf{p}(T_{i,-}), \mathbf{lv}(T_{i,-}), \mathbf{er}(T_i)]$;

$\mathbf{info}'(T_i) \leftarrow \text{LinearLayer}(\mathbf{info}(T_i))$;

$i \leftarrow 0$;

**for** $mask$ $in$ $\mathbf{m}(T_i)$ **do**

  **if** $mask = 0$ **then**

    ▷ Here $v[i:j]$ $(j > i)$ denotes the $i$-th element to the $j$-th element (not including $j$) of $v$, for any vector $v$. $H$ denotes hidden size

    $\mathbf{info}'(T_i)[i \times H + 1 : (i+1) \times H + 1] \leftarrow 0$;

  **end**

  $i \leftarrow i + 1$;

**end**

▷ In this context, "GRUCell" refers to a single GRU cell defined in [1]

$\mathbf{h}(T_{i,+}) \leftarrow \text{GRUCell}(\mathbf{info}'(T_i), \mathbf{h}(T_{i,-}))$;

$\mathbf{p}(T_{i,+}) \leftarrow g_p(\mathbf{h}(T_{i,+}))$;

$\mathbf{lv}(T_{i,+}) \leftarrow g_{lv}(\mathbf{h}(T_{i,+}))$;

### 3 Structure of $g_p$ and $g_{lv}$

The computation of $g_p$ and $g_{lv}$ is performed using a single neural network, allowing simultaneous calculation of both. The hidden state $\mathbf{h}$, which encodes the system's dynamics, is initially passed through a fully connected linear layer that transforms it into a space matching the number of prediction units. To introduce nonlinearity, a Rectified Linear Unit (ReLU) activation function is applied, followed by a dropout layer to enhance generalization by randomly deactivating a fraction of the activations during training. The resulting intermediate representation is then processed through another fully connected linear layer, which outputs a vector of size $2 \times$ input size. A sigmoid activation function is applied to this output to ensure that all values are constrained within a biologically meaningful range of 0 to 1. The final output is then split into two parts: the first half represents the predicted mean ($\mathbf{p}$), while the second half corresponds to the log variance ($\mathbf{lv}$).

### 4 Univariate simulation settings

#### 4.1 Settings for different groups

- **Group 1:** Utilizes the Black-Scholes process with parameters $k = 0.8$, $\sigma = 0.2$, and initial values ranging from 1 to 2. The time series generated follows a positive trend.

- **Group 2:** Employs the Black-Scholes process with parameters $k = -0.5$, $\sigma = 0.2$, and initial values ranging from 15 to 18, resulting in a time series with a negative trend.

- **Group 3:** Implements the Ornstein-Uhlenbeck process with parameters $k = 5$, $m = 20$, $\sigma = 4$, and initial values between 2 and 3, generating a time series that reverts to a mean of 20.

- **Group 4:** Uses the Ornstein-Uhlenbeck process with parameters $k = 5$, $m = 4$, $\sigma = 1.5$, and initial values ranging from 15 to 18, producing a time series that reverts to a mean of 4.
- **Group 5:** Applies the Ornstein-Uhlenbeck process with parameters $k = 5$, $m = 10$, $\sigma = 1.5$, and initial values between 5 and 18, creating a time series that reverts to a mean of 10.

##### 4.2 Settings for different scenarios

In the first simulation scenario, $\pi$ is set to 0.15, indicating a 15% chance of data being undetectable, and data points are randomly dropped with a probability of $\delta = 0.15$, resulting in 15% of the detectable data being unavailable. Therefore, the remaining information $\eta$ is approximately $0.85 \times 0.85 \approx 72\%$. For the second simulation, we set both $\pi$ and $\delta$ to 0.25, corresponding to a proportion of remaining information $\eta$ being approximately $0.75 \times 0.75 \approx 56\%$. Similarly, in the third simulation, the values of $\pi$ and $\delta$ are increased to 0.35, resulting in $\eta$ being approximately $0.65 \times 0.65 \approx 42\%$; and in the fourth simulation, $\pi$ and $\delta$ are further increased to 0.4, leading to $\eta$ of approximately $0.6 \times 0.6 \approx 36\%$.

#### 5 Multivariate simulation settings

##### 5.1 Settings for different groups

- **Group 1:** $\lambda_1 = 0.8$, $\lambda_2 = 0.1$ for this group.
  1. **Microbiome 1:** Utilizes the Ornstein-Uhlenbeck process with parameters $k = 5$, $m = 20$, $\sigma = 4$, and initial values ranging from 2 to 3. The time series generated tends to revert to a mean level of 20.
  2. **Microbiome 2:** Utilizes the Black-Scholes process with parameters $k = 1.5$, $\sigma = 0.2$, and initial values ranging from 1 to 2. The time series generated follows a positive trend.
  3. **Microbiome 3:** Utilizes the Ornstein-Uhlenbeck process with parameters $k = 5$, $m = 4$, $\sigma = 1.5$, and initial values ranging from 15 to 18. The time series generated tends to revert to a mean level of 4.
  4. **Microbiome 4:** Utilizes the Black-Scholes process with parameters $k = -0.5$, $\sigma = 0.2$, and initial values ranging from 15 to 18. The time series generated follows a negative trend.
  5. **Microbiome 5:** Utilizes the Ornstein-Uhlenbeck process with parameters $k = 5$, $m = 10$, $\sigma = 1.5$, and initial values ranging from 5 to 18. The time series generated tends to revert to a mean level of 10.
- **Group 2:** $\lambda_1 = 0.8$, $\lambda_2 = 0.1$ for this group.
  1. **Microbiome 1:** Utilizes the Black-Scholes process with parameters $k = 0.9$, $\sigma = 0.25$, and initial values ranging from 1 to 3. The time series generated follows a positive trend.
  2. **Microbiome 2:** Utilizes the Ornstein-Uhlenbeck process with parameters $k = 4$, $m = 15$, $\sigma = 2$, and initial values ranging from 10 to 12. The time series generated tends to revert to a mean level of 15.
  3. **Microbiome 3:** Utilizes the Black-Scholes process with parameters $k = -0.7$, $\sigma = 0.15$, and initial values ranging from 5 to 7. The time series generated follows a negative trend.
  4. **Microbiome 4:** Utilizes the Ornstein-Uhlenbeck process with parameters $k = 6$, $m = 8$, $\sigma = 1.2$, and initial values ranging from 2 to 4. The time series generated tends to revert to a mean level of 8.
  5. **Microbiome 5:** Utilizes the Black-Scholes process with parameters $k = 1.2$, $\sigma = 0.3$, and initial values ranging from 3 to 5. The time series generated follows a positive trend.

- **Group 3:** $\lambda_1 = 0.8$, $\lambda_2 = 0.03$ for this group.
  1. **Microbiome 1:** Utilizes the Ornstein-Uhlenbeck process with parameters $k = 3$, $m = 12$, $\sigma = 2.5$, and initial values ranging from 4 to 6. The time series generated tends to revert to a mean level of 12.
  2. **Microbiome 2:** Utilizes the Black-Scholes process with parameters $k = 0.5$, $\sigma = 0.1$, and initial values ranging from 2 to 4. The time series generated follows a positive trend.
  3. **Microbiome 3:** Utilizes the Ornstein-Uhlenbeck process with parameters $k = 7$, $m = 5$, $\sigma = 1$, and initial values ranging from 8 to 10. The time series generated tends to revert to a mean level of 5.
  4. **Microbiome 4:** Utilizes the Black-Scholes process with parameters $k = -1$, $\sigma = 0.2$, and initial values ranging from 10 to 12. The time series generated follows a negative trend.
  5. **Microbiome 5:** Utilizes the Ornstein-Uhlenbeck process with parameters $k = 5$, $m = 9$, $\sigma = 1.5$, and initial values ranging from 1 to 3. The time series generated tends to revert to a mean level of 9.
- **Group 4:** $\lambda_1 = 0.8$, $\lambda_2 = 0.03$ for this group.
  1. **Microbiome 1:** Utilizes the Black-Scholes process with parameters $k = 1$, $\sigma = 0.25$, and initial values ranging from 3 to 5. The time series generated follows a positive trend.
  2. **Microbiome 2:** Utilizes the Ornstein-Uhlenbeck process with parameters $k = 4$, $m = 7$, $\sigma = 2$, and initial values ranging from 5 to 7. The time series generated tends to revert to a mean level of 7.
  3. **Microbiome 3:** Utilizes the Black-Scholes process with parameters $k = -0.8$, $\sigma = 0.15$, and initial values ranging from 7 to 9. The time series generated follows a negative trend.
  4. **Microbiome 4:** Utilizes the Ornstein-Uhlenbeck process with parameters $k = 6$, $m = 10$, $\sigma = 1.2$, and initial values ranging from 2 to 4. The time series generated tends to revert to a mean level of 10.
  5. **Microbiome 5:** Utilizes the Black-Scholes process with parameters $k = 1.1$, $\sigma = 0.3$, and initial values ranging from 4 to 6. The time series generated follows a positive trend.
- **Group 5:** $\lambda_1 = 0.8$, $\lambda_2 = 0.03$ for this group.
  1. **Microbiome 1:** Utilizes the Ornstein-Uhlenbeck process with parameters $k = 3$, $m = 11$, $\sigma = 2.5$, and initial values ranging from 3 to 5. The time series generated tends to revert to a mean level of 11.
  2. **Microbiome 2:** Utilizes the Black-Scholes process with parameters $k = 0.6$, $\sigma = 0.1$, and initial values ranging from 1 to 3. The time series generated follows a positive trend.
  3. **Microbiome 3:** Utilizes the Ornstein-Uhlenbeck process with parameters $k = 7$, $m = 6$, $\sigma = 1$, and initial values ranging from 6 to 8. The time series generated tends to revert to a mean level of 6.
  4. **Microbiome 4:** Utilizes the Black-Scholes process with parameters $k = -1.2$, $\sigma = 0.2$, and initial values ranging from 8 to 10. The time series generated follows a negative trend.
  5. **Microbiome 5:** Utilizes the Ornstein-Uhlenbeck process with parameters $k = 5$, $m = 8$, $\sigma = 1.5$, and initial values ranging from 2 to 4. The time series generated tends to revert to a mean level of 8.

#### 5.2 Settings for different scenarios

In the first simulation, $\alpha$ is set to $-5$, $\beta$ is set to $-0.5$, and data points are randomly dropped with a probability of $\delta = 0.12$. The remaining information $\eta$ is approximately 70%. In the second simulation, $\alpha$ is set to $-5$, $\beta$ is set to 0, and data points are randomly dropped with a probability of $\delta = 0.3$. The remaining information $\eta$ is approximately 50%. In the third simulation, $\alpha$ is set to $-5$, $\beta$ is set to 1, and data points are randomly dropped with a probability of $\delta = 0.39$. The remaining information $\eta$ is approximately 30%. In the fourth simulation, $\alpha$ is set to $-5$, $\beta$ is set to 1.5, and data points are randomly dropped with a probability of $\delta = 0.48$. The remaining information $\eta$ is approximately 20%.

### 6 Species grouping in ECC dataset

To reduce redundancy and enhance interpretability, certain species in the ECC dataset are combined into groups based on established taxonomic classifications. Below, we detail the composition of each group:

- **Mitis Group:** This group includes *Streptococcus australis*, *Streptococcus cristatus* (formerly *Streptococcus crista*), *Streptococcus dentisani*, *Streptococcus gordonii*, *Streptococcus infantis*, *Streptococcus mitis*, *Streptococcus oligofermentans*, *Streptococcus oralis*, *Streptococcus parasanguinis* (formerly *Streptococcus parasanguis*), *Streptococcus peroris*, *Streptococcus pneumoniae*, *Streptococcus pseudopneumoniae*, *Streptococcus sanguinis* (formerly *Streptococcus sanguis*), and *Streptococcus tigurinus*.
- **Salivarius Group:** This group includes *Streptococcus salivarius*, *Streptococcus vestibularis*, and *Streptococcus thermophilus*.
- **Streptococcus Anginosus Group (The Milleri Group):** This group includes *Streptococcus constellatus*, *Streptococcus intermedius*, *Streptococcus anginosus*, and *Streptococcus sinensis*.
- **Group A Streptococcus:** This group includes *Streptococcus pyogenes*, commonly known as Group A strep.
- **Mutans Streptococci Group (The S. Mutans Group):** This group includes *Streptococcus mutans*, *Streptococcus sobrinus*, and *Streptococcus downei*.

### 7 Additional figures

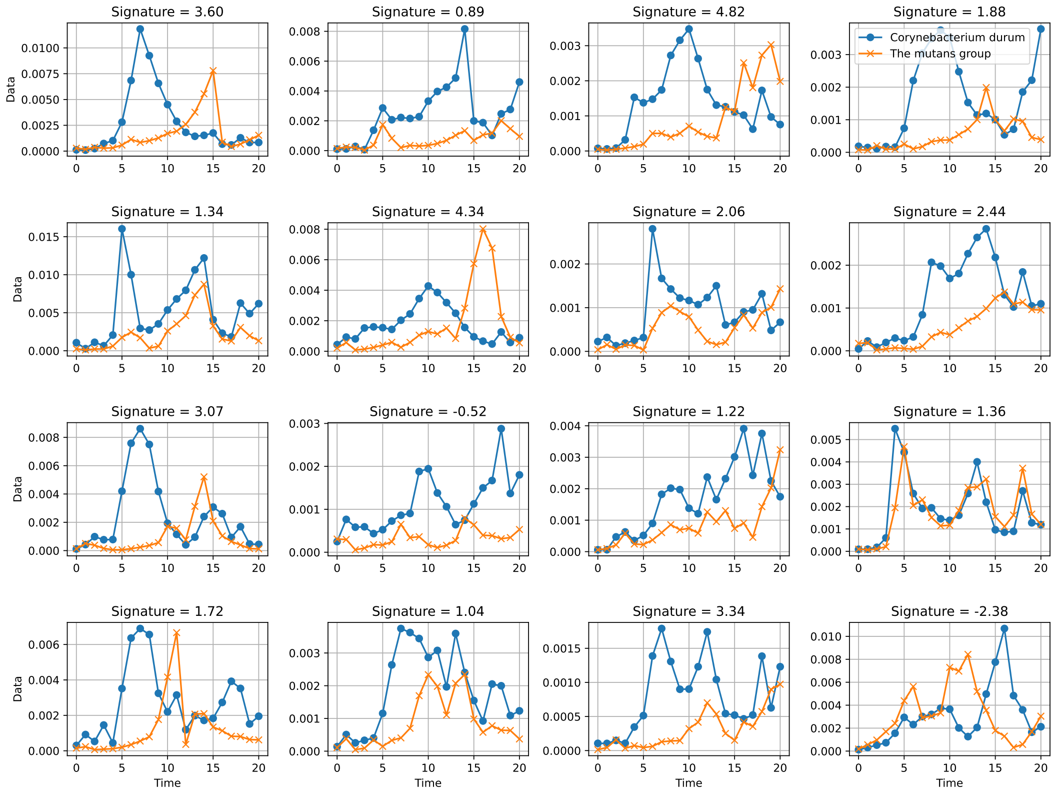


**Figure 1.** The lead-lag relationship and the computed pairwise signatures of of interpolated Corynebacterium durum and the mutans group in 16 representative subjects for ECC. y-axis indicates relative abundance of a species, x-axis indicates time interval. Higher signature indicates *Corynebacterium durum* has a stronger lead effect on *mutans*.

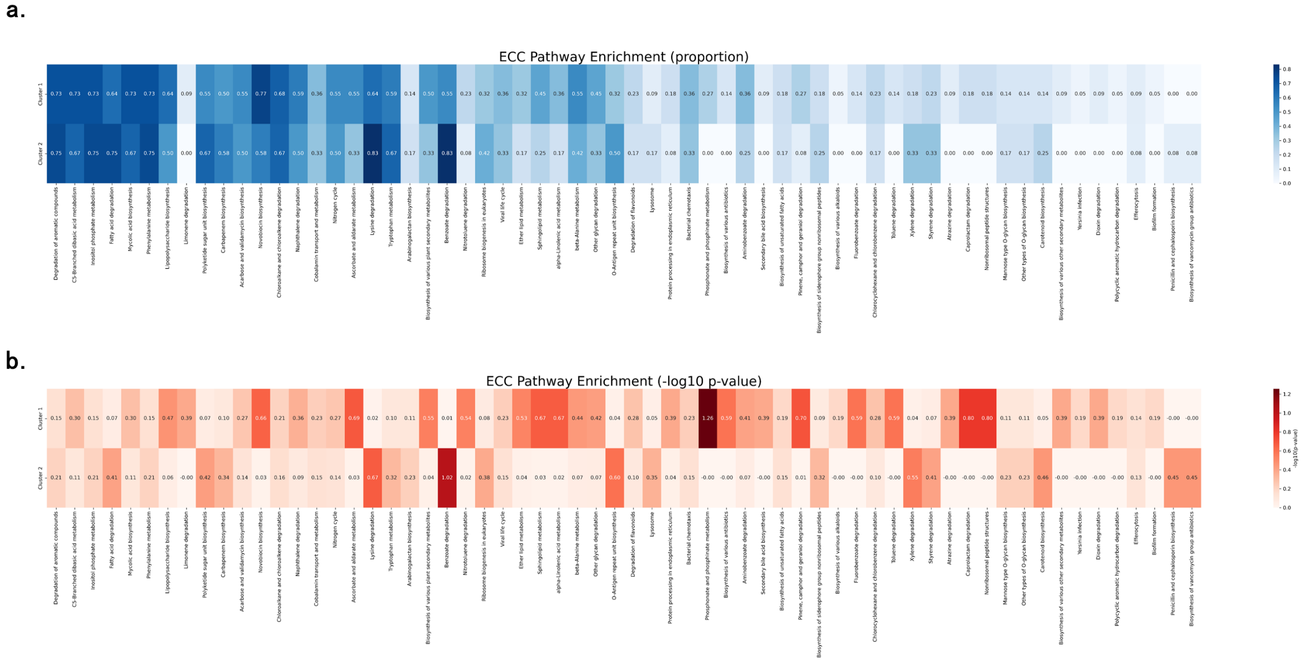


**Figure 2.** **(a)** Proportion of microbes within clusters involved in the pathway in the ECC dataset. **(b)** -log10(p-value) for each pathway in each cluster (Fischer's Exact Test).

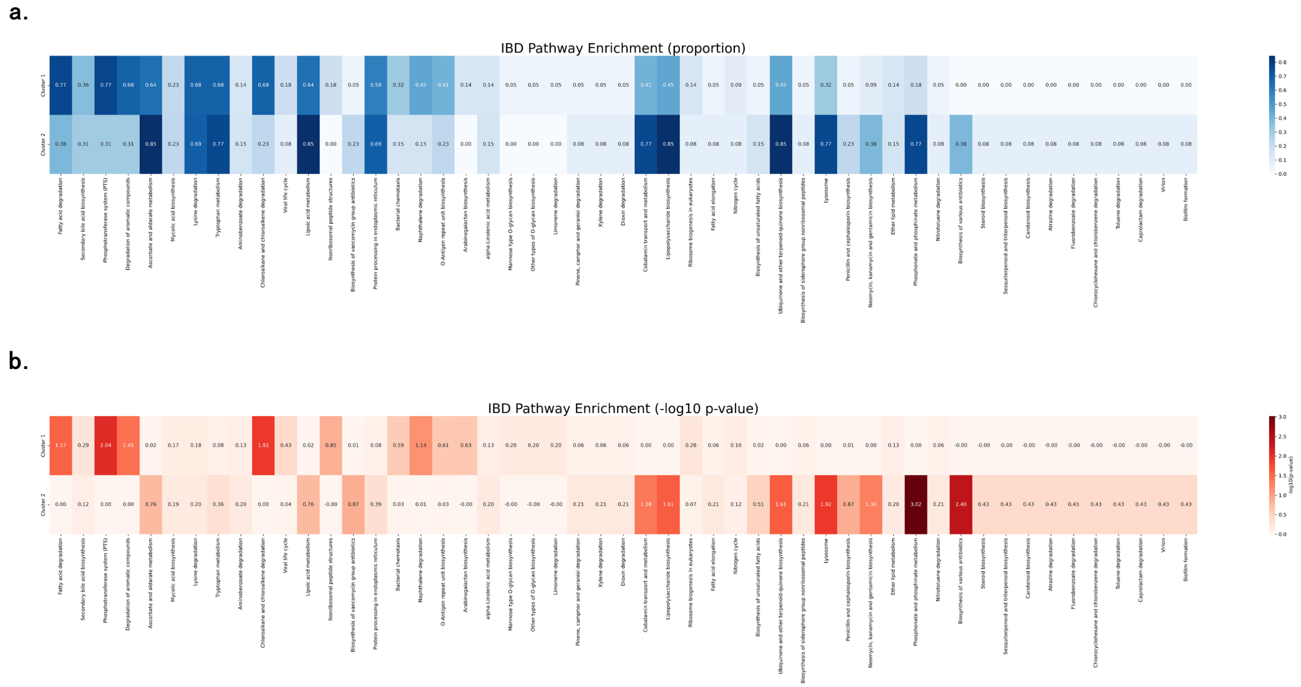


**Figure 3. (a)** Proportion of microbes within clusters involved in the pathway in the IBD dataset. **(b)** -log10(p-value) for each pathway in each cluster (Fischer's Exact Test).
